# Mechanistic Plasticity of the RabGEFs Mon1-Ccz1 and Fuzzy-Inturned

**DOI:** 10.1101/2025.03.27.645700

**Authors:** Stephan Wilmes, Jesse Tönjes, Jan-Hannes Schäfer, Anna Lürick, Dovile Januliene, Steven Apelt, Daniele Di Iorio, Seraphine V Wegner, Martin Loose, Arne Moeller, Daniel Kümmel

## Abstract

Rab GTPases organize intracellular trafficking and provide identity to organelles. Their spatiotemporal activation by guanine nucleotide exchange factors (GEFs) is tightly controlled to ensure fidelity. Our structural and functional comparison of the tri-longin domain RabGEFs Mon1-Ccz1 and Fuzzy-Inturned reveals the molecular basis for their target specificity. Both complexes rely on a conserved sequence motif of their substrate GTPases for the catalytic mechanism, while secondary interactions allow discrimination between targets. We also find that dimeric Mon1-Ccz1 from fungi and the metazoan homologs with the additional third subunit RMC1/Bulli bind membranes through electrostatic interactions via distinct interfaces. Protein-lipid interaction studies thus reveal a function of RMC1/Bulli in the complex as mediator of membrane recruitment. In the case of Fuzzy-Inturned, reconstitution experiments demonstrate that the BAR (Bin-Amphiphysin-Rvs) domain protein CiBAR1 can support membrane recruitment of the GEF. Collectively, our study demonstrates the molecular basis for the adaptation of TLD-RabGEFs to different cellular functions.

## Introduction

Organization of the eucaryotic endomembrane system critically depends on the precise spatiotemporal control of Rab GTPases (*1*). As markers of organelle identity and part of the conserved fusion machinery, Rabs are required for communication and exchange between different intracellular membrane compartments (*2*, *3*). They cycle between an active GTP-bound form that associates with membranes via prenyl anchors and an inactive GDP-bound form that is kept cytosolic through association with a GDI (guanosine dissociation inhibitor) chaperone. Depending on the nucleotide loading state, two switch regions of the Rab GTPase adopt different conformations that interact with different binding partners. The switch between both states is intrinsically slow and tightly controlled by regulatory proteins. GAPs (GTPase activating proteins) stimulate GTP hydrolysis resulting in inactivation. Activation requires GEFs (guanine nucleotide exchange factors) that reduce the nucleotide affinity of the Rab GTPase and thus allow loading with GTP. Because the GTP-bound Rab can no longer bind to GDI, activation of Rabs simultaneously leads to its membrane recruitment. The localization of RabGEFs therefore also determines the subcellular distribution of their cognate Rab GTPases and represents a key event in achieving specificity in intracellular trafficking.

The tri-longin domain (TLD) RabGEF complexes represent a family of RabGEFs with a unique conserved structural core (*4*, *5*). In each complex, two subunits that contain three longin domains (LD1-3) each form a heterodimer, which is required for catalytic activity towards the cognate Rab GTPases. Different TLD-GEFs vary in their composition as they can contain additional domains or bind auxiliary subunits (Fig. 1A).

**Fig. 1:**
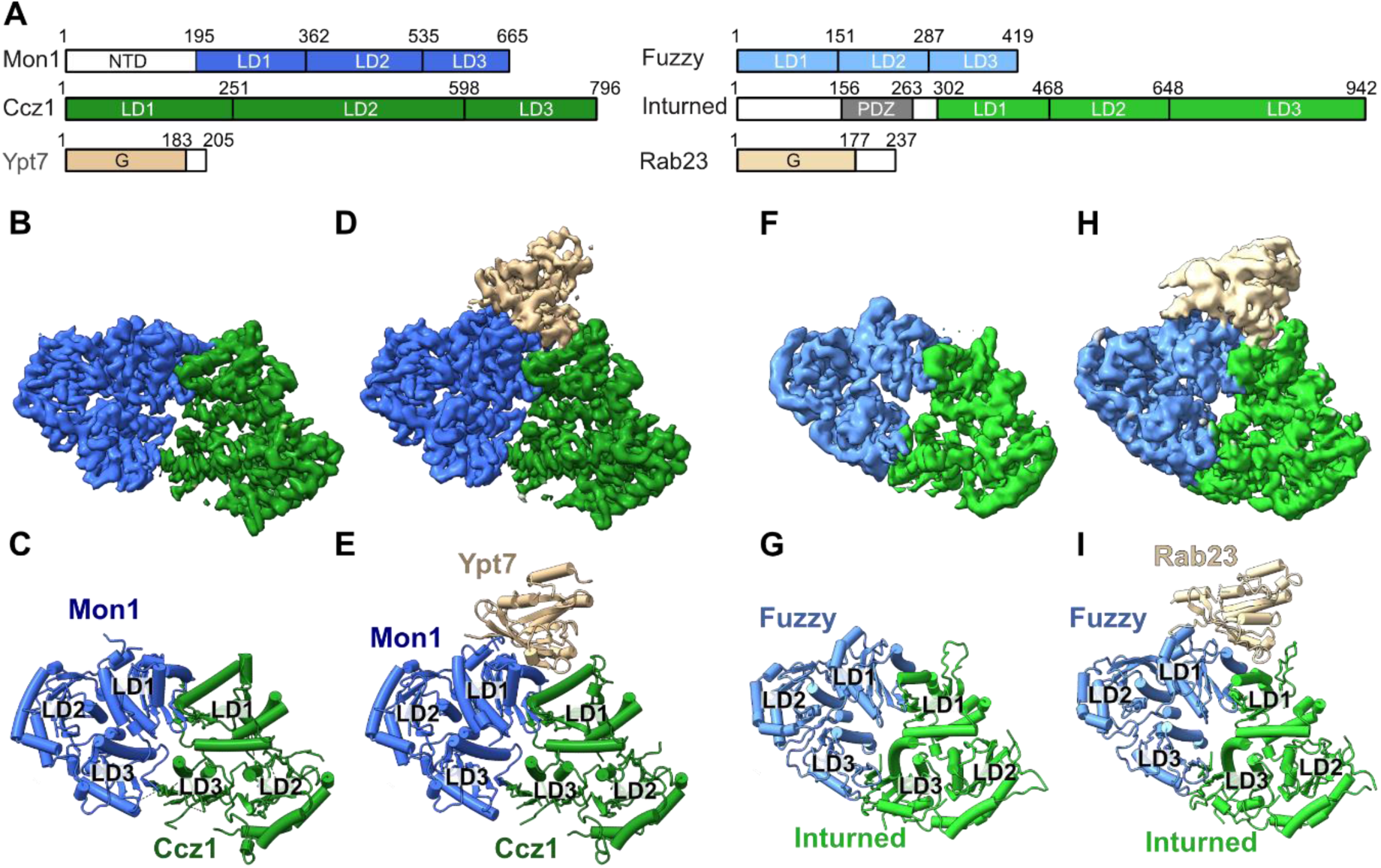
Structural analysis of TLD GEF-Rab complexes. (**A**) Domain architecture of *Ct*Mon1, *Ct*Ccz1, *Ct*Ypt7, *Hs*Fuzzy, *Hs*Inturned, and *Hs*Rab23. LD: longin domain, NTD: N-terminal domain, G: GTPase domain, PDZ: PSD95/Dlg1/ZO-1 domain. (**B**) Cryo-EM map and (**C**) cartoon representation of the dimeric *Ct*Mon1-Ccz1^ΔL^ complex. (**D**) Cryo-EM map and (**E**) cartoon representation of the trimeric *C*tMon1-Ccz1^ΔL^-Ypt7^N125I^ complex. (**F**) Cryo-EM map and (**G**) cartoon representation of the dimeric *Hs*Fuzzy-Inturned complex. (**H**) Cryo-EM map and (**I**) cartoon representation of the trimeric *Hs*Fuzzy-Inturned-Rab23^N121I^ complex.

The Mon1-Ccz1 complex is the best-studied TLD GEF and conserved throughout all eukaryotes (*6–10*). Mon1-Ccz1 is a dimer in yeast and activates the Rab7 homolog Ypt7. This process is required in endosomal maturation and autophagy, where generation of Ypt7-positive compartments are destined for fusion with the vacuole, the lysosome-equivalent organelle of yeast (*11*). Structural and functional studies have provided a framework for a mechanistic understanding of Mon1-Ccz1 (*12–15*). The first longin domains of Mon1 and Ccz1 form the active site and bind Ypt7 to remodel switch 1 of the GTPase (*14*). As a consequence, a conserved lysine is inserted into the nucleotide-binding pocket and expels the essential magnesium cofactor, leading to lowered nucleotide affinity. The localization of Mon1-Ccz1 to endosomes requires binding to active Rab5-like GTPases and binding to phosphatidylinositol phosphate (PIP). Autophagosomal recruitment, however, relies on the interaction of Mon1-Ccz1 with Atg8 and with lipid packing defects. Thus, Mon1-Ccz1 is differentially targeted by distinct molecular cues (*16*). In metazoans, Mon1-Ccz1 form a trimer with an additional subunit Bulli/RMC1 and control Rab7 activation, fulfilling a conserved function in organizing lysosomal fusion (*7*, *8*, *10*). While Bulli/RMC1 has been shown to be functionally important *in vivo*, its molecular role remains elusive.

The BLOC-3 (biogenesis of lysosome-related organelles complex-3) complex, composed of the TLD subunits Hps1 and Hps4, is unique to metazoans and activates Rab32 and Rab38 for the maturation of lysosome-related organelles (*17*). Interestingly, mutations in the HPS1 and HPS4 genes cause the genetic disease Hermansky-Pudlak syndrome, but the molecular mechanisms underlying BLOC-3 function are poorly understood (*18*).

Finally, the TLD proteins Fuzzy and Inturned act as GEF for Rab23 (*5*). In a trimeric complex with the protein Fritz, Fuzzy-Inturned constitute the PPE (planar cell polarity effector) complex in *Drosophila melanogaster* (*Dm*) and are involved in establishing planar cell polarity (*19*). In mammalian cells, Inturned, Fuzzy, the Fritz homolog Wdpcp and the atypical GTPase Rsg1 form the CPLANE (ciliogenesis and planar cell polarity effector) complex that is required for cilia formation (*20*). The molecular understanding of Fuzzy and Inturned function remains incomplete. A reported cryo-EM structure of CPLANE established structural similarity between Fuzzy-Inturned and Mon1-Ccz1 (*21*). Furthermore, binding to PIP lipids has been suggested for several CPLANE subunits but it is not clear how the complex is regulated and localized in cells.

We here report the cryo-EM structures of the Mon1-Ccz1 and Inturned-Fuzzy core complexes bound to their cognate GTPases Ypt7 and Rab23 in the nucleotide exchange transition states, revealing a conserved catalytic mechanism with distinct modes of substrate recognition. Furthermore, we quantitatively show that fungal Mon1-Ccz1 robustly interacts with lipids, whereas metazoan Mon1-Ccz1 and Inturned-Fuzzy complexes do not. Instead, metazoan TLD-GEFs depend on binding to accessory subunits and recruiter proteins. For CPLANE, we demonstrate that the BAR (Bin-Amphiphysin-Rvs) domain adaptor protein CiBAR1/FAM92A1 (*22–24*) can fulfill this role. These findings advance our understanding of Rab regulation by TLD RabGEFs in metazoans, which is relevant fundamental physiological processes like ciliogenesis and related pathological scenarios.

## Results

### Structural analysis of substrate-bound Mon1-Ccz1 and Fuzzy-Inturned

TLD RabGEFs have been proposed to utilize a conserved catalytic mechanism but reportedly maintain a high specificity for their respective substrate GTPases (*5*, *14*). To provide the molecular basis of these observations, we determined the cryo-EM structures of fungal Mon1-Ccz1 from *Chaetomium thermophilum* (*Ct*) bound to nucleotide-free Ypt7 and of human Fuzzy-Inturned in complex with nucleotide-free Rab23.

In our previous work, we resolved a crystal structure of a catalytic *Ct*Mon1^LD1^-Ccz1^LD1^-Ypt7^N125I^ subcomplex (*14*) and the cryo-EM structure of a truncated *Ct*Mon1^ΔN^-Ccz1^ΔL^ complex lacking the N-terminal domain of Mon1 (NTD, residues 1-140) and the lipid-binding loop of Ccz-1 (L, residues 333-444) (*12*). However, the N-terminal region of Mon1 was shown to play a role in regulating the complex activity (*25*). Furthermore, it has been shown that some Rab GEF complexes like TRAPP (*26*) or Rig1-Rgp1 (*27*) have high affinity binding sites for the hypervariable domain (HVD) of their substrate GTPases. Recognition of secondary motifs in addition to G domain binding has emerged as an important factor for Rab specificity (*26–29*). To investigate potential interactions of Mon1^NTD^ and Ypt7^HVD^ with the TLD GEF core complex, we use full-length Mon1 and Ypt7 in the current study. To support the formation of a stable transition state of the complex, the nucleotide-free variant *Ct*Ypt7^N125I^ was co-expressed with *Ct*Mon1 and *Ct*Ccz1^ΔL^ in *E. coli*. Analysis of purified complexes by cryo-EM revealed a mixture of dimeric *Ct*Mon1-Ccz1 and trimeric *Ct*Mon1-Ccz1-Ypt7 particles. We could determine the structure of both complexes at 3 Å resolution (Fig. 1B-E, S1, S2, S3A,B). The structure of the dimeric complex with full-length Mon1 closely resembles the structure of truncated Mon1^ΔN^-Ccz1^ΔL^ (r.m.s.d. 1.17 Å over 837 residues, Fig. S3C), but the higher resolution of the current structure allowed us to improve the sequence assignment of the final β-strand of Ccz1. In the trimeric complex, the structures of Mon1 and Ccz1 remain virtually unchanged (r.m.s.d. 0.67 Å, Fig. 1), and we assigned the additional density to the G domain of Ypt7. We did not observe structural changes in binding of Ypt7 to intact Mon1-Ccz1 in comparison to the Mon1^LD1^-Ccz1^LD1^ subcomplex (r.m.s.d. 1.03 Å over 436 residues). The experimental map does not reveal unmodelled density that could be interpreted as the Mon1^NTD^ or Ypt7^HVD^. Consequently, our structures do not provide support for models that suggest an autoinhibitory interaction of the NTD of Mon1 with the longin domains or recognition of the Ypt7^HVD^ by Mon1-Ccz1 at the chosen conditions. However, these interactions may require the presence of membranes or recruiter proteins.

To resolve the *Hs*Fuzzy-Inturned-Rab23 complex, we again used a nucleotide-free mutant of the GTPase (Rab23^N121I^) and performed co-expression in insect cells. From the analysis of the cryo-EM data set, we could determine the structure of the dimeric Fuzzy-Inturned complex in addition to the trimeric *Hs*Fuzzy-Inturned-Rab23^N121I^ complex at 3.6 Å and 3.4 Å resolution, respectively (Fig. 1F-I, S4, S5, S6A,B). The conformations of Fuzzy and Inturned do not change upon Rab23 binding (r.m.s.d. 0.82 Å, Fig. 1) and are virtually identical compared to the full CPLANE complex that also includes Rsg1 and Wdpcp (r.m.s.d. 1.25 Å over 748 residues, Fig. S6C) (*21*). All longin domains of Fuzzy and Inturned are well resolved, but as in CPLANE, an N-terminal PDZ domain of Inturned is not visible in the map.

The GTPases interact in a very similar orientation with same interfaces of the GEF complexes, which are provided by the first longin domains of the TLD subunits (Fig. 1E,I, 2A,B). Fuzzy and Mon1 fulfill homologous functions and contribute the majority of the interface with Rab23 (885 Å^2^) and Ypt7 (1270 Å^2^), respectively. However, Inturned and Ccz1 also both make critical contacts with the switch 1 regions (428 Å^2^ and 469 Å^2^, respectively). Overall, the structures of Mon1-Ccz1-Ypt7 and Inturned-Fuzzy-Rab23 complexes show a highly similar architecture (r.m.s.d. 2.59 Å over 747 aligned residues).

### Conserved and variable features of the TLD RabGEF mechanism

Remodeling of the Ypt7 switch 1 region by Mon1-Ccz1 causes an opening of the nucleotide binding pocket, removal of a guanine base interacting phenylalanine (F33) and the insertion of a catalytic lysine residue (K38) in the binding site that expels the magnesium cofactor (Fig. 2A). F33 and a tyrosine (Y37) adjacent to the catalytic lysine bind to hydrophobic pockets on Mon1-Ccz1, thus stabilizing the switch 1 conformation. Collectively, this structural arrangement was shown to lower the nucleotide affinity of Ypt7 and thus promote nucleotide exchange (*14*).

**Fig. 2:**
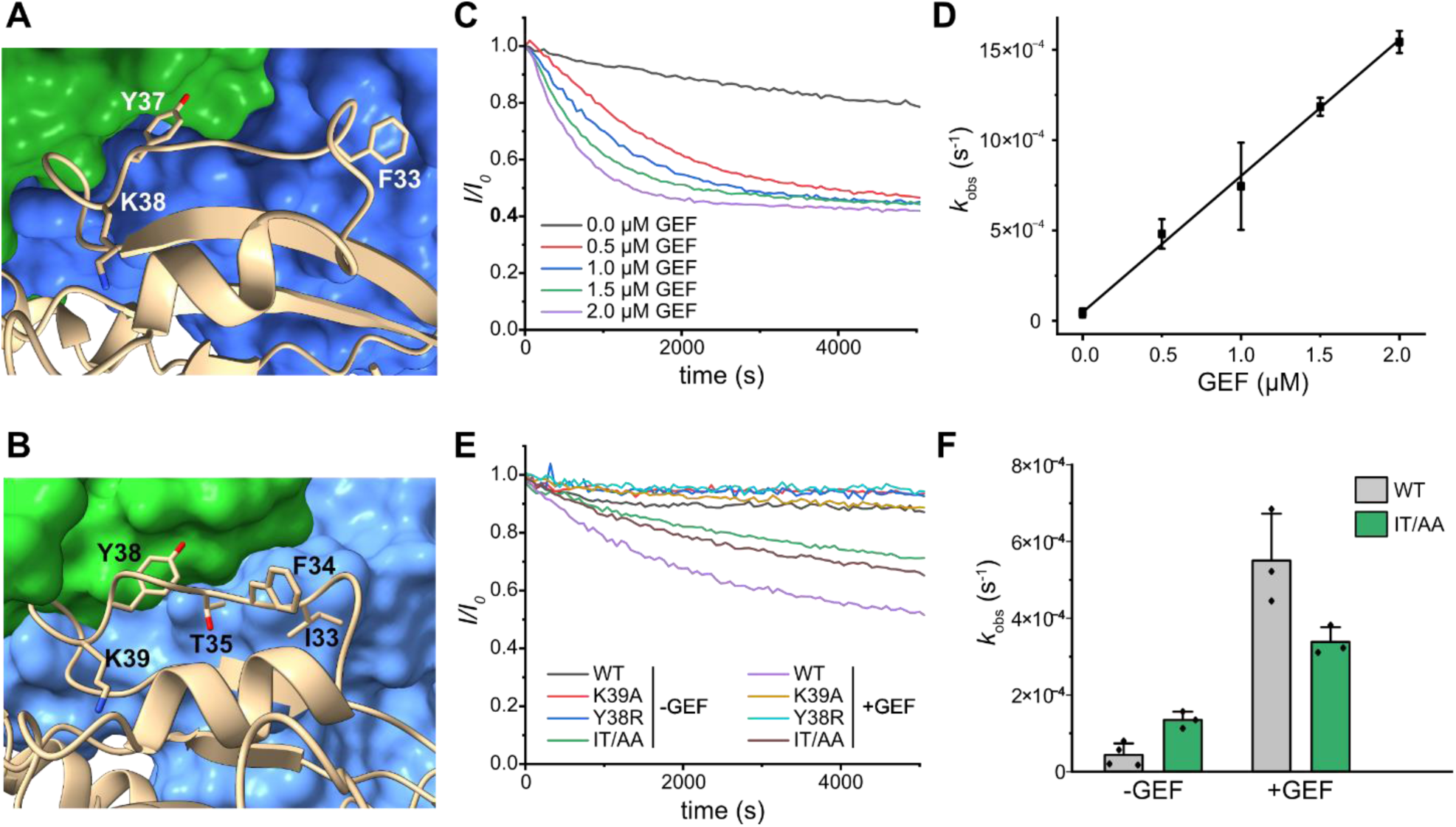
Catalytic GEF mechanism. (**A**) Close-up of the Ypt7 switch 1 region (tan) interacting with Mon1 (blue) and Ccz1 (green). (**B**) Close-up of the Rab23 switch 1 region (wheat) interacting with Fuzzy (blue) and Inturned (green). (**C**) Guanine nucleotide exchange assay of Rab23 with different concentrations of Fuzzy^LD1^-Inturned^LD1^ catalytic core complex. (**D**) Determination of catalytic efficiency from n=3 independent experiments as in C. Data are represented as means ± standard deviation. (**E**) Guanine nucleotide exchange assay of Rab23 wild-type (WT) and different variants (IT/AA: I33A, T35A) with and without Fuzzy^LD1^-Inturned^LD1^. (**F**) Intrinsic and GEF-stimulated nucleotide exchange rates of Rab23 WT and the IT/AA variant from n=3 repeats. Data are represented as means ± standard deviation.

A basic residue at the position of the catalytic lysine is conserved in the Inturned-Fuzzy substrate Rab23 (K39) and the BLOC-3 substrates Rab32 and Rab38 (R55 and R38, respectively), as well as the adjacent tyrosine (Fig. S7A). This suggested that the catalytic mechanism of TLD RabGEFs is likely conserved. Indeed, when bound to its GEF, K39 and Y38 of Rab23 adopt the same conformation as the corresponding residues of Ypt7 (Fig. 2B). We used an *in vitro* GEF assay with an Fuzzy^LD1^-Inturned^LD1^ core complex, which showed robust nucleotide exchange activity on wt Rab23 (k_cat_/K_M_ of 7×10^2^ M^-1^ sec^-1^, Fig. 2C,D). However, upon mutation of either of the conserved residues K39 to alanine or Y38 to arginine, Rab23 cannot be stimulated by Fuzzy-Inturned (Fig. 2E). Thus, the common YK motif in TLD RabGEF substrates is part of a conserved mechanism. The switch 1 region of Rab23 is well resolved in the cryo-EM map (Fig. S6B), and variations in the GEF interaction compared to Ypt7 can be observed that likely encode substrate specificity. The guanine base stabilizing aromatic residue F34 of Rab23 is removed from the binding pocket, but in contrast to F33 of Ypt7, it does not bind to a hydrophobic pocket on the GEF surface. Instead, residues I33 and T35 interact with Inturned and thus fix the switch 1 conformation (Fig. 2B). A double mutant Rab23^I33A/T35A^ showed increased intrinsic nucleotide exchange but reduced stimulation of nucleotide exchange by Fuzzy-Inturned (Fig. 2E,F). Because structures of Rab23 in the GTP- and GDP-bound states (*30*) showed that I33 and T35 do not interact with the nucleotide directly (Fig. S7B,C), the increased intrinsic exchange rate is likely caused by a destabilization of switch 1. However, the ∼5-fold diminished ability of Fuzzy-Inturned to promote nucleotide exchange of Rab23^I33A/T35A^ confirms the importance of these amino acids for recognition by the GEF and for the catalytic function. Thus, TLD RabGEFs rely on both conservation and variability in the switch 1 regions of their substrates to realize catalytic efficiency and specificity.

### Membrane binding of metazoan Mon1-Ccz1 requires Bulli

In addition to substrate recognition, spatiotemporal control of RabGEF localization has been recognized as a key mechanism for their specificity and regulation (*1*, *28*). The analysis of the electrostatic surface potential of Mon1-Ccz1 revealed a large basic patch on the Mon1^LD2^ located opposite of the Ypt7 binding site that - together with an amphipathic helix in a disordered region of Ccz1^LD2^ (*16*) - mediates binding of Mon1-Ccz1 to model membranes (Fig. 3A). For a quantitative determination of membrane affinity, we turned to QCM-D (quartz crystal microbalance with dissipation monitoring) measurements. Supported lipid bilayer (SLBs) were formed with a DO/PIP/PS lipid mix that displays packing defects and negative charges (Fig. S8A). Here, the titration of *Ct*Mon1-Ccz1 on the SLBs yielded an apparent *K*_D_ of 67 nM (Fig. 3B,C).

**Fig. 3:**
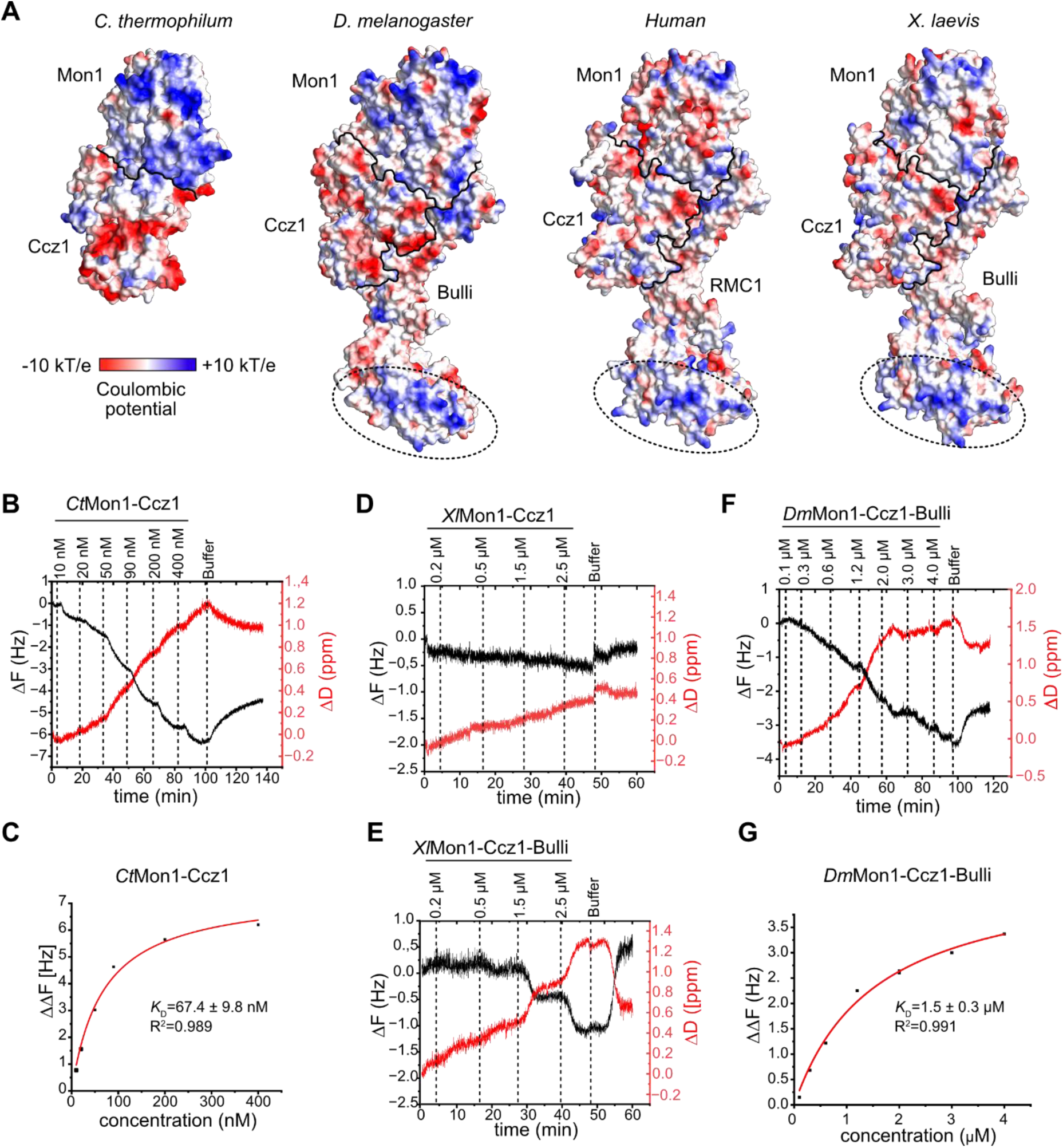
Membrane binding of fungal and metazoan Mon1-Ccz1 complexes. (**A**) Surface representation of the lipid interaction side *Ct*Mon1-Ccz1 and of *Dm*Mon1-Ccz1-Bulli, *Hs*Mon1-Ccz1-RMC1 and *Xl*Mon1-Ccz1-Bulli in the corresponding orientation. Coloring is by electrostatic potential. Subunit boundaries are indicated by black lines and a conserved basic patch on Bulli/RMC1 is highlighted by circles. (**B**) QCM-D measurements of a *Ct*Mon1-Ccz1 titration on a supported lipid bilayer (74 mol% DO-PC, 18 mol% DO-PE, 2 mol% DO-PI3P, 1 mol% DO-PI3,5P_2_, 5 mol% DO-PS). (**C**) QCM-D frequency shifts of *Ct*Mon1-Ccz1 plotted against the concentration of protein were fitted with a one-step binding model for *K*_D_ determination. (**D**) QCM-D measurements of a *Xl*Mon1-Ccz1 titration on a supported lipid bilayer as in B. (**E**) QCM-D measurements of a *Xl*Mon1-Ccz1-Bulli titration on a supported lipid bilayer as in B. (**F**) QCM-D measurements of a *Dm*Mon1-Ccz1-Bulli titration on a supported lipid bilayer as in B. (**G**) QCM-D frequency shifts of *Dm*Mon1-Ccz1-Bulli plotted against the concentration of protein were fitted with a one-step binding model for *K*_D_ determination.

The importance of lipid recognition via a basic patch and amphipathic helix in conjunction with recruiter proteins for the function of Mon1-Ccz1 in fungi has been established *in vitro* and in cellular studies with yeast (*11*, *16*, *31*). To assess if the localization mechanism of the trimeric metazoan Mon1-Ccz1 containing complexes may be conserved, we investigated the experimental structure of *Dm*Mon1-Ccz1-Bulli (*13*, *15*) and AlphaFold models (*32*) of human (*Hs*) Mon1-Ccz1-RMC1 and *Xenopus laevis* (*Xl*) Mon1-Ccz1-Bulli (Fig. S8B-G). The analysis of the coulomb potential of the membrane binding interface of the different metazoan complexes reveals that the prominent basic patch of Mon1 is not conserved (Fig. 3A), suggesting that lipid interactions of these complexes may be weaker than those of *Ct*Mon1-Ccz1. Of note, clusters of basic residues are located on the β-propeller of Bulli/RMC1 (Fig. 3A), which interestingly are conserved (*13*). We next asked to what extent direct lipid binding as a recruitment mechanism can also be observed for the metazoan Rab7 GEF homologs. With QCM-D, we could not detect membrane binding of dimeric *Xl*Mon1-Ccz1, but of trimeric *Xl*Mon1-Ccz1-Bulli (Fig. 3D,E). Because the membrane binding of *Xl*Mon1-Ccz1-Bulli is weak and the protein was not stable at concentrations > 2.5 µM needed to reach saturation in a titration experiment, we could not calculate the *K*_D_ of the interaction. We detected a similarly weak interaction of *Dm*Mon1-Ccz1-Bulli with supported bilayer by QCM-D (Fig. 3F). For the trimeric complex from flies, we could calculate a *K*_D_ of ∼1.5 µM, which is almost 25 times weaker than for *Ct*Mon1-Ccz1. Taken together, these results demonstrate a conserved membrane-binding behavior of metazoan Mon1-Ccz1-Bulli/RMC1 complexes that requires the auxiliary third subunit Bulli/RMC1.

### Membrane recruitment of Fuzzy-Inturned by a BAR-domain protein adaptor

The membrane targeting mechanism of Fuzzy and Inturned has not been studied in detail but both proteins have been reported to interact with monophosphorylated PIPs - with a preference for PI3P - in protein-lipid overlay assays (*21*). However, analysis of the surface properties of the Fuzzy-Inturned complex reveals a mixed distribution of positive and negative charges and no conserved basic or hydrophobic patches. (Fig. 4A, S9). We therefore tested the membrane binding of Fuzzy-Inturned in liposome sedimentation assays. Fuzzy-Inturned does bind neither liposomes generated from a neutral lipid mix that has few packing defects (PO) nor charged liposomes with increased packing defects (DO/PIP/PS) (Fig. 4B). We used Mon1-Ccz1 as a control (*12*, *16*), which does not associate with neutral liposomes but is robustly recruited to charged liposomes with packing defects (Fig. S10A). We also could not detect any interaction of Fuzzy-Inturned with liposomes that have a high content of 5% PI3P (Fig. 4C). Our findings are not consistent with the previously suggested function of Inturned and Fuzzy as PIP-binding proteins (*21*), pointing to alternative mechanisms.

**Fig. 4:**
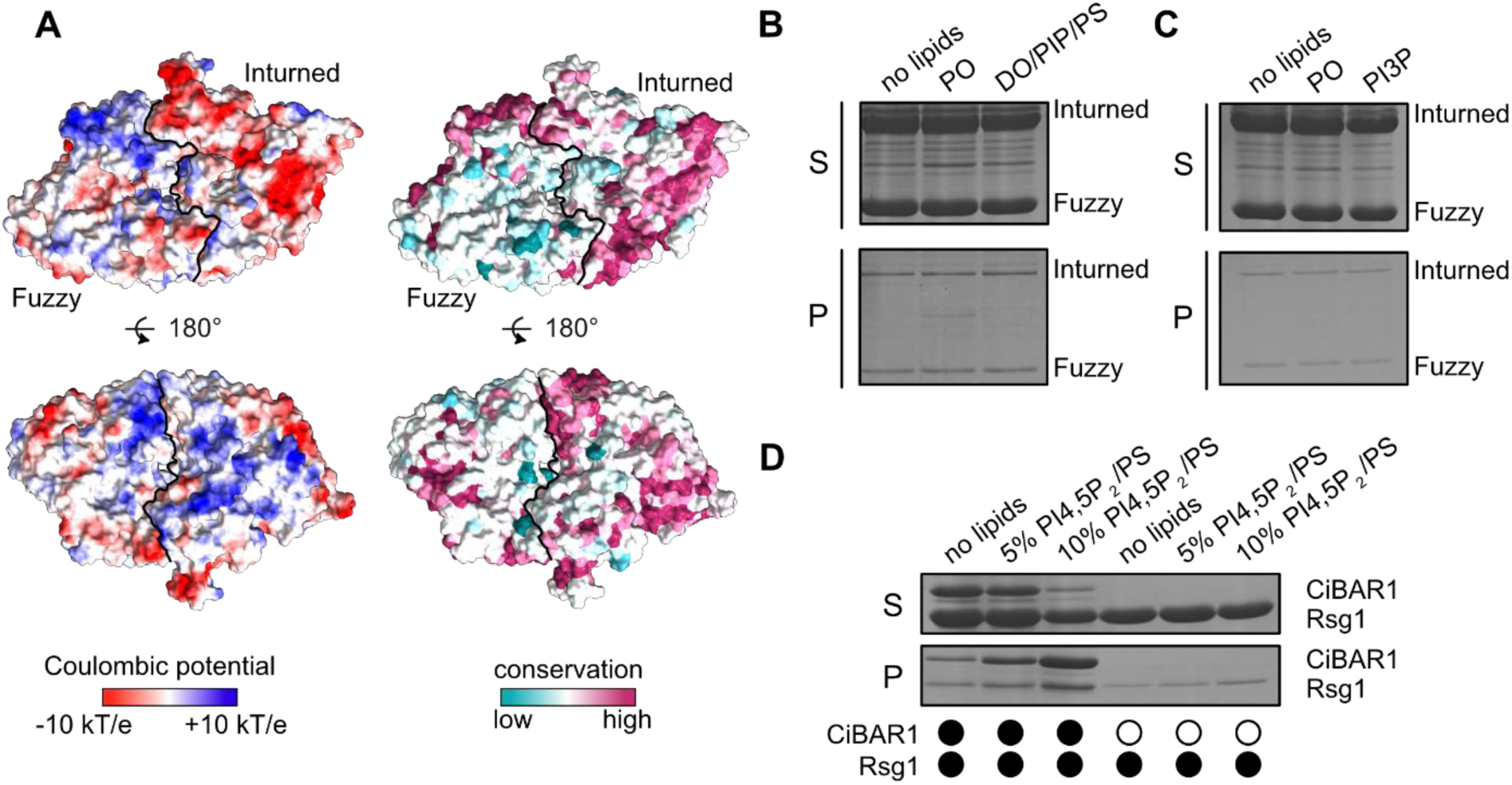
Membrane recruitment of Fuzzy-Inturned. (**A**) Surface representation of Fuzzy-Inturned colored by electrostatic potential or conservation in two opposing views. (**B**) Sedimentation assay of Fuzzy-Inturned with no, neutral PO (81 mol% PO-PC, 18 mol% PO-PE, 1 mol% DP-PE-Atto565), and charged DO/PIP/PS (73 mol% DO-PC, 18 mol% DO-PE, 1 mol% DP-PE-Atto565, 2 mol% PI3P, 1 mol% PI3,5P_2_, 5 mol% PS) liposomes. (**C**) Sedimentation assay of Fuzzy-Inturned with no, neutral PO (81 mol% PO-PC, 18 mol% PO-PE, 1 mol% DP-PE-Atto565), and liposomes with high PI3P (76 mol% PO-PC, 18 mol% PO-PE, 1 mol% DP-PE-Atto565, 5 mol% PI3P). (**D**) Sedimentation assay of Rsg1 and a CiBAR1-Rsg1 complex with liposomes containing varying amounts of PI4,5P_2_. Sedimentation assays were analyzed by SDS-PAGE and Coomassie staining.

Recruiter proteins play a key role in the localization of RabGEF complexes (*11*, *25*, *33*). From the notion that RabGEFs are frequently recruited by upstream GTPases in trafficking pathways the concept of Rab cascades has emerged (*34*). In the family of TLD GEFs the GTPase Rab5 is described as recruiter for Mon1-Ccz1(-Bulli) (*13*, *35*) and Rab9 for the BLOC-3 complexes (*36*). Thus, the Fuzzy-Inturned complex may also be localized to its target compartments by interactions with recruiter proteins that enable tight membrane binding. It is therefore tempting to speculate that the GTPase Rsg1, which binds Fuzzy-Inturned (*20*), may serve as recruiter for this complex. However, Rsg1 does not contain a lipidation site or any other membrane anchoring motif. Consistent with this notion, we also did not observe a direct interaction of Rsg1 with liposomes containing packing defects and PIPs in sedimentation assays (Fig. 4D).

Interestingly, a recent study reported an interaction of Rsg1 with the classical BAR domain protein CiBAR1 (*24*), which forms a curved helix dimer and interacts with charged lipids (Fig. S10B) (*22*, *23*). Thus, CiBAR1 may be an adaptor that links Fuzzy-Inturned via Rsg1 to membranes. We co-expressed CiBAR1 and Rsg1 and were able to purify a complex of both proteins with equimolar stoichiometry as judged from a Coomassie stained SDS-PAGE gel (Fig. S10C). The recombinant CiBAR1-Rsg1 complex was efficiently recruited to PI4,5P_2_ containing liposomes (Fig. 4D), consistent with the PIP-specificity reported for CiBAR1 before (*23*). Thus, CiBAR1 fulfills the biochemical properties to serve as an adaptor protein for the mammalian CPLANE complex.

Based on these findings, we created a model of a CiBAR1-bound CPLANE complex using AlphaFold3 (Fig. S10D-F) (*32*). All subunits Rsg1, Fuzzy, Inturned and Wdpcp are predicted to bind above or at the putative membrane interaction interface as defined by the concave PIP-interaction surface of CiBAR1 (Fig. 5A). In this model, the β-propeller of Wdpcp would align at the membrane with a conserved basic patch (Fig. 5B). This is reminiscent of our findings regarding Bulli and suggests that both proteins, although their structural integration in the complex is entirely different (Fig. S11), may fulfill a similar function in orienting the complexes on the membrane. Because CiBAR1 is a symmetric dimer that can bind two molecules of Rsg1, we docked two CPLANE complexes to the adaptor protein, which could be accommodated without major clashes (Fig. 5C).

**Fig. 5:**
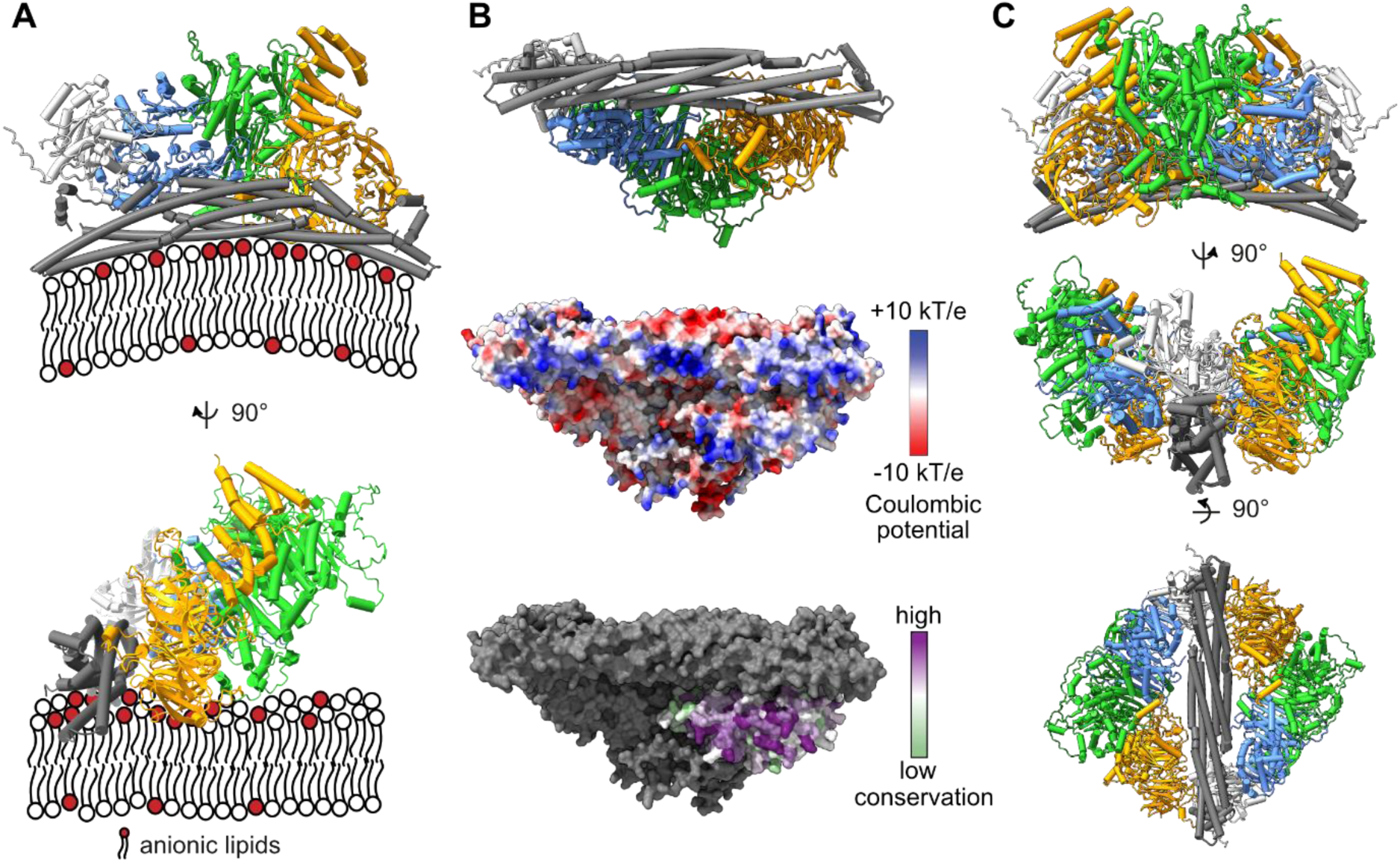
Model for CPLANE membrane recruitment. (**A**) AlphaFold3 prediction of a CPLANE (Rsg1 (light grey) - Fuzzy (blue) – Inturned (green) – Wdpcp (orange))-CiBAR1 (drak grey) complex in two perpendicular orientations positioned on a lipid bilayer as defined by the membrane interaction interface of CiBAR1. (**B**) Membrane-facing view of CPLANE-CiBAR1 in cartoon representation, as surface representation with electrostatic potential and as surface representation with Wdpcp colored by conservation. (**C**) Model of a decameric complex containing two CPLANE complexes (colored as in A) docked onto a CiBAR1 dimer.

## Discussion

Our comparative study of Mon1-Ccz1 and Fuzzy-Inturned complexes reveals that the catalytic mechanism of these RabGEFs requires a Y-K motif in their substrate GTPases. The related BLOC-3 activates GTPases with a Y and an R at the equivalent position (*17*), which defines a Y-K/R consensus motif as a requirement for TLD RabGEF substrates. Interestingly, we identified four additional Rab GTPases with this motif at the correct position in their switch 1 region (Fig. S5C). Because the GEFs for some of these Rabs are not known, they might represent novel targets for TLD RabGEFs. However, in the case of Rab7L, which has such a Y-K motif, it was already shown that it is not a substrate of any of the known TLD RabGEFs (*5*).

With respect to membrane recruitment, we observe a striking difference between the fungal Mon1-Ccz1 complexes, which strongly bind to charged lipids, and the metazoan Mon1-Czz1-Bulli homologs, which only bind weakly. Also, while the electrostatic interactions of the fungal complexes are primarily mediated by the Mon1 subunit (*37*), metazoan complexes depend on Bulli/RMC1 for membrane association (Fig. 3). These findings provide the first demonstration of a molecular function of these additional subunits only found in metazoans. Loss of Bulli/RMC1 dramatically affects functionality of the endolysosomal system (*7*, *8*, *10*), but the catalytic GEF function of Mon1-Ccz1 is largely retained in the absence of a third subunit (*8*). A supporting role in membrane recruitment is therefore a likely function of Bulli/RMC1, which could explain the observed phenotypes and adds another layer of spatiotemporal control during endosomal maturation.

Our work does not confirm a PIP-specificity of Fuzzy and Inturned, as reported in a previous study that used protein-lipid overlay assays (*21*). The preferential interaction with PI3P may represent a false positive resulting from the caveats of this assay where lipids are spotted on a nitrocellulose membrane and are not incorporated into a lipid bilayer. Because PIP strips represent the less physiological setup, we would tentatively rely on the results from sedimentation assays and conclude that Fuzzy and Inturned do not specifically recognize PI3P.

In addition to direct lipid interactions, membrane targeting of Mon1-Ccz1 depends on the binding to lipid anchored recruiter proteins. Associations with Atg8 or Rab5 GTPases via an effector interaction contribute to autophagosomal or endosomal localization, respectively (*11*, *16*, *33*). For the metazoan Mon1-Ccz1-Bulli/RMC1 complexes, with reduced membrane affinity compared to the fungal homologs, the activity of the GEF and membrane targeting is dominated by recruiter proteins (*25*). It is therefore plausible that Fuzzy-Inturned will be regulated similarly.

Our findings establish CiBAR1 as a recruiter protein for CPLANE. This protein has been linked to ciliogenesis and is a likely mediator for directing Fuzzy-Inturned activity and Rab23 to the basal body. Interestingly, CiBAR1 binds Chibby1 (*38*), which in turn interacts with the Rab8 GEF Rabin8 (*39*). Thus, it is tempting to speculate that the activation of two key GTPases for the progression of ciliogenesis, Rab8 and Rab23, may be coordinated.

Fuzzy-Inturned is linked to its membrane recruiter protein via the small GTPase Rsg1. The role of Rsg1 in ciliogenesis and as part of CPLANE is established (*20*, *40*, *41*), but little is known about its molecular function. While Rsg1 engages Fuzzy in an effector mode in its GTP-bound form (*21*), the computational model indicates that the interaction with CiBAR is nucleotide-independent. This contrasts with other examples of BAR domain interactions with Rho or Rab GTPases, which require the active, GTP-loaded GTPases (*42*, *43*). It will be interesting to investigate if a GEF and/or GAP for Rsg1 exist that may regulate its nucleotide loading state. The interaction between CiBAR1 and Rsg1 should thereby not be affected, but the binding to Fuzzy and thus the membrane targeting of CPLANE could potentially be regulated.

Because Fuzzy-Inturned is not only implicated in ciliogenesis, it is possible that additional adaptors besides CiBAR1 can direct Fuzzy-Inturned to other locations. For example, the recruitment of the complex in planar cell polarity signaling may involve different mechanisms. The identity of the required factors involved in this pathway remains to be investigated.

Our proposed model of a fully assembled membrane-associated CiBAR1-CPLANE complex predicts that a conserved basic patch of the subunit Fritz may support membrane binding. This is reminiscent of our findings regarding Bulli, which is a structural homolog of Fritz. The structural integration of Fritz and Bulli within their respective complexes is entirely different (Fig. S11), which could point to an adaptation to accommodate a similar function in orienting the complexes on the membrane. This observation further supports the main conclusions of our study, which reveal mechanisms of molecular fine-tuning in TLD family GEF complexes to achieve specialization for dedicated functions.

## Materials and Methods

### Protein Expression and Purification

#### HsFuzzy-Inturned-Rab23^N121I^

The complex of human FLAG-Inturned, His-SUMO-Fuzzy and nucleotide free GST-Rab23^N121I^ was produced in *Spodoptera frugiperda* 21 cells using the biGBac expression system (*44*). Cells were lysed in buffer A (100 mM Tris pH 8, 200 mM NaCl, 0.5 mM TCEP, 1 mM MgCl_2_) supplemented with 0.25% Igepal CA-630, 0.025 mg/ml DNaseI and protease inhibitor mix HP (Serva). Complexes were isolated on glutathione agarose and eluted by proteolytic cleavage of the GST affinity tag. Proteins were further purified via size exclusion chromatography (SEC650 10/300, Bio-Rad) with buffer B (50 mM Tris, 200 mM NaCl, 0.5 mM TCEP, and 1 mM MgCl_2_, pH 8).

#### CtMon1-Ccz1^ΔL^-Ypt7^N125I^

*E. coli* Bl21(DE3) cells were co-transformed with GST-PreScission-Mon1, His6-SUMO-Ccz1^ΔL^ (lacking residues 360-460), and nucleotide free GST-TEV-Ypt7^N125I^ and expression (16 h, 16 °C) induced after 30 minute cold shock by 0.25 mM isopropyl-β-D-thiogalactoside. Cells were lysed in buffer C (50 mM NaH_2_PO_4_, 500 mM NaCl, 1 mM MgCl_2_, 5% (v/v) glycerol, pH 7.3) supplemented with protease inhibitor mix HP (Serva), 0.025 mg/ml DNase I and 1 mg/ml Lysozyme. Cleared lysates were incubated with glutathione Agarose, the resin washed and affinity tags cleaved off Mon1 and Ccz1. After additional washing steps, the complex was eluted by TEV cleavage of the GST tag on Ypt7^N125I^. Proteins were further purified by size exclusion chromatography (SEC650 10/300, Bio-Rad) with buffer D (25 mM HEPES, 250 mM NaCl, 1 mM MgCl_2_, 0.5 mM TCEP, pH 7.3)

#### Rab23 variants

*E. coli* Bl21(DE3) cells were transformed with GST-Rab23 variants (WT, K39A, Y38R, I33A/T35A) cloned into pCDF6P vectors and expression (16 h, 16 °C) induced after 30 minute cold shock by 0.25 mM isopropyl-β-D-thiogalactoside. Cells were lysed in buffer C (500 mM NaCl, 50 mM NaH_2_PO_4_, 5% Glycerol, 2 mM DTT, 1 mM MgCl_2_ pH 7.5) supplemented with protease inhibitor mix HP (Serva), 0.025 mg/ml DNase I and 1 mg/ml Lysozyme. Cleared lysates were incubated with glutathione Agarose, the resin washed and proteins eluted by addition of 20 mM glutathione and 10 mM DTT. Proteins were further purified by size exclusion chromatography (SEC650 10/300, Bio-Rad) with buffer D (200 mM NaCl, 10 mM HEPES, 1 mM MgCl_2_, 5% Glycerol, 0.5 mM TCEP, pH 7.3)

#### HsFuzzy-Inturned LD1 dimer

*E. coli* Bl21(DE3) cells were co-transformed with GST-Inturned^LD1^ and His6-SUMO-Fuzzy^LD1^ and expression (16 h, 16 °C) induced after 30 minute cold shock by 0.25 mM isopropyl-β-D-thiogalactoside. Cells were lysed in buffer C (500 mM NaCl, 50 mM NaH_2_PO_4_, 5% Glycerol, 2 mM DTT, 1 mM MgCl_2_ pH 7.5) supplemented with protease inhibitor mix HP (Serva), 0.025 mg/ml DNase I and 1 mg/ml Lysozyme. Cleared lysates were incubated with glutathione Agarose, the resin washed and proteins eluted by addition of 20 mM glutathione and 10 mM DTT. Proteins were further purified by size exclusion chromatography (SEC650 10/300, Bio-Rad) with buffer D (200 mM NaCl, 10 mM HEPES, 1 mM MgCl_2_, 5% Glycerol, 0.5 mM TCEP, pH 7.3)

#### CtMon1-Ccz1

*E. coli* Bl21(DE3) cells were transformed with pCDF6P *Ct*Mon1 and pET28HS *Ct*Ccz1 and expression (16 h, 16 °C) induced after 30 minute cold shock by 0.25 mM isopropyl-β-D-thiogalactoside. Cells were lysed in buffer C (500 mM NaCl, 50 mM NaH_2_PO_4_, 5% glycerol, 2 mM DTT, 1 mM MgCl_2_ pH 7.5) supplemented with protease inhibitor mix HP (Serva), 0.025 mg/ml DNase I and 1 mg/ml Lysozyme. Cleared lysates were incubated with glutathione Agarose, the resin washed and His-Sumo tag proteolytically cleaved off Ccz1. The resin was further washed and the protein complex finally eluted by removal of the GST-tag of Mon1 through incubation with PreScission protease. Proteins were further purified by size exclusion chromatography (SEC650 10/300, Bio-Rad) with buffer E (250 mM NaCl, 25 mM HEPES, 1 mM MgCl_2_, 0.5 mM TCEP, pH 7.3)

#### XlMon1-Ccz1-Bulli and XlMon1-Ccz1

Complexes were produced in *Spodoptera frugiperda* 21 cells using the biGBac expression system (*44*). Cells were lysed in buffer F (50 mM HEPES-KOH, pH 7.8, 0.5 mM EDTA, 150 mM KOAc, 1 mM DTT, 10% Glycerol) supplemented with 0.1% NP40, 0.025 mg/ml DNaseI and protease inhibitor mix HP (Serva). Complexes were isolated on a streptactin matrix and eluted with 2.5 mM d-desthiobiotin. Affinity tags were cleaved by TEV protease incubation and proteins further purified via size exclusion chromatography (SEC650 10/300, Bio-Rad) with buffer G (25 mM HEPES-NaOH, pH 7.3, 250 mM NaCl, 1 mM MgCl_2_, 0.5 mM TCEP).

#### HsRsg1

A template plasmid encoding *Hs*Rsg1 was a kind gift from Francis Barr, University of Oxford. *E. coli* Bl21(DE3) cells were transformed with pCDF6P hsRsg1 and expression (16 h, 16 °C) induced after 30 minute cold shock by 0.25 mM isopropyl-β-D-thiogalactoside. Cells were lysed in buffer H (200 mM NaCl, 20 mM HEPES pH 8, 1 mM DTT, 1 mM MgCl_2_) supplemented with protease inhibitor mix HP (Serva), 0.025 mg/ml DNase I and 1 mg/ml Lysozyme. Cleared lysates were incubated with glutathione Agarose, the resin washed and proteins eluted by incubation with PreScission.

#### HsCiBAR1-Rsg1

Flag-CiBAR1 was a gift from Ken-Ichi Takemaru (Addgene plasmid # 200440) (*38*). The CiBAR1 insert was subcloned into a pET28HS vector with a His6-SUMO tag. *E. coli* Rosetta (DE3) pLysS cells were co-transformed with pET28HS-CiBAR1 and pCDF6P-Rsg1 constructs and expression (16 h, 16 °C) was induced after a 30 minute cold shock by addition of 0.25 mM isopropyl-β-D-thiogalactoside. Cells were lysed in buffer I (150 mM NaCl, 20 mM HEPES, 1 mM DTT, 1 mM MgCl_2_ pH 7.4) supplemented with protease inhibitor mix HP (Serva), 0.025 mg/ml DNase I and 1 mg/ml Lysozyme. Cleared lysates were incubated with glutathione agarose, the resin was washed and the His-Sumo tag was proteolytically cleaved off CiBAR1. The resin was further washed and the protein complex was eluted by removal from the GST-tag of Rsg1 through incubation with PreScission protease.

#### *In vitro* GEF Assay

Guanine nucleotide exchange assays were performed by loading purified GTPases with 1.5 M excess MANT-GDP in the presence of 20 mM EDTA for 30 min at 30 °C. After quenching the loading reaction with 25 mM MgCl_2_, the Rab23-MANT-GDP complex was separated from excess MANT-GDP in Nap5 columns (Cytiva). GTPases (2 µM final concentration) and the minimal GEF complex (varying concentrations between 0 and 2 µM) were mixed in buffer J (200 mM NaCl,10 mM HEPES, 1 mM MgCl_2_, 0.5 mM TCEP, pH 7.3) and nucleotide exchange reaction started by adding 0.1 mM GTP. The reaction was monitored in microplate reader (M1000 Pro, Tecan) tracing the decrease of fluorescence at λ_em_ 448 nm (λ_ex_ 354 nm) in intervals of 30 s or 60 s at 25 °C. Data were fitted with a first-order exponential decay function y=y_0_ + A. e^−x/t^ to calculate *k_obs_* = τ^−1^ (s^−1^) using OriginPro (OriginLab). Catalytic efficiency was determined by plotting *k_obs_* against the concentration of the GEF.

### Co-Sedimentation Assay

Lipids were mixed in chloroform and dried in a SpeedVac The lipid film was dissolved in buffer L (25 mM HEPES, 250 mM NaCl, 1 mM MgCl_2_, pH 7.3) supplemented with 5% sucrose to a final lipid concentration of 2 mM. Multilamellar lipid vesicles were generated by five cycles of freezing in liquid nitrogen and thawing at 56 °C and stored at −80 °C. Prior to use, multilamellar vesicles were extruded 21 times through a polycarbonate membrane to generate liposomes of 400 nm diameter.

Proteins and liposomes were mixed in buffer L in final concentrations of 1 μM (Fuzzy-Inturned, Mon1-Ccz1), 2 µM (CiBAR1 and Rsg1) and 0.5 mM (lipids), respectively. Reactions were incubated for 20 min at room temperature, and liposomes were pelleted at 20,000 x *g* for 20 min at 4 °C. The supernatant fraction was precipitated with acetone at −20 °C, and supernatant and pellet fractions were analyzed by SDS-PAGE and Coomassie staining.

### Membrane compositions

<u>PO:</u> 81 mol% PO-PC, 18 mol% PO-PE, 1 mol% DP-PE-Atto565

<u>DO/PIP/PS:</u> 73 mol% DO-PC, 18 mol% DO-PE, 1 mol% DP-PE-Atto565, 2 mol% PI3P, 1 mol% PI3,5P_2_, 5 mol% PS

<u>PI3P:</u> 76 mol% PO-PC, 18 mol% PO-PE, 1 mol% DP-PE-Atto565, 5 mol% PI3P

5% PI4,5P_2_/PS: 71 mol% PO-PC, 18 mol% PO-PE, 1 mol% DP-PE-Atto565, 5 mol% PI4,5P_2_, 5 mol% PS

10% PI4,5P_2_/PS: 61 mol% PO-PC, 18 mol% PO-PE, 1 mol% DP-PE-Atto565, 10 mol% PI4,5P_2_, 10 mol% PS

### Quartz crystal microbalance with dissipation (QCM-D)

QCM-D measurements were performed essentially as previously described (*45*, *46*). Multilamellar vesicles were freshly extruded to 100 nm through a polycarbonate membrane. QCM-D measurements were performed with a QSense Analyser (Biolin scientific, Gothenburg, Sweden) and SiO_2_-coated sensors (QSX 303, 50 nm SiO2, 4.95 MHz) equipped with four temperature-controlled flow cells. Measurements were performed at a working temperature of 23°C with a peristaltic pump (Ismatec, Grevenbroich, Germany) with a flow rate of 75 µl/min for supported lipid bilayer (SLB) formation and 35 µl/min for protein binding. The sensors were activated by a 11 minute treatment with a UV/ozone cleaner (Ossila, Sheffield, UK) prior to SLB formation. For SLB formation, freshly prepared SUVs were diluted to 0.1 mg/ml with citrate buffer (10 mM tri-sodium citrate, 150 mM NaCl, 10 mM CaCl_2_ pH 4.6) and flushed into the chamber. For protein binding, reaction buffer (25 mM HEPES-NaOH, 250 mM NaCl, 1 mM MgCl_2_, 0.5 mM TCEP pH 7.3) was used. Proteins were titrated in increasing concentrations until a stable baseline was reached. The change in frequency (ΔΔF) of the fifth overtone resonance frequency channel was plotted against the protein concentration. Nonlinear curve fitting assuming one site specific binding was performed in OriginPro (OriginLab) to determine the *K*_D_.

### Cryo-EM sample Preparation

Purified *Ct*Mon1-Ccz1^ΔL^-Ypt7^N125I^ (∼0.8 mg/ml) and *Hs*Fuzzy-Inturned-Rab23^N121I^ (∼0.7 mg/ml) complexes were applied to glow-discharged CF-1.2/1.3-3 Cu-50 grids. 0.002% lauryl maltose neopentyl glycol (LMNG) was added to *Hs*Fuzzy-Inturned-Rab23^N121I^ on grid prior to freezing to overcome preferred orientation of particles at the air water interface. Samples were plunge frozen in liquid ethane using a Vitrobot Mark IV (ThermoFisher Scientific). Data were collected on a Glacios electron microscope equipped with a Selectris energy filter and a Falcon 4i direct electron detector (ThermoFisher Scientific) at 165,000-fold nominal magnification.

### Cryo-EM Image Processing

All cryo-EM data were preprocessed in cryoSPARC Live, and further processing was performed in cryoSPARC v3 and v4 (Fig. S1 and S3) (*47–49*).

#### CtMon1-Ccz1^ΔL^-Ypt7^N125I^

After micrograph curation, 25142 micrographs with a CTF fit better than 5 Å were included for further data analysis (Fig. S1). Initial particles were selected by blob picker implemented in CryoSPARC Live, and additional particle picking was performed using cryoSPARC’s Topaz wrapper (*50*). After particle duplicate removal and particle extraction in a box size of 400 pixels (Fourier-cropped to 200 pixels, resulting in a pixel size of 1.36 Å per pixel), iterative rounds of 2D classification and selection were performed for all particle stacks obtained from each of the picking jobs to eliminate bad picks (Fig. S1). After 2D classifications, ab-initio reconstruction with six classes followed by heterogeneous refinement was performed. The heterogeneous refinement produced two classes, revealing trimeric (Mon1-Ccz1-Ypt7) or dimeric (Mon-Ccz1) particle projections. These classes were combined in Non-uniform (NU) refinements and submitted to subsequent iterative 3D classification with forced hard classification enabled, which resulted in two good classes containing either the dimeric or the trimeric complexes. These two classes (525 thousand and 1.27 million particles) were further refined using NU refinement in a box size of 440 pixels (without Fourier cropping). The obtained reconstructions of both populations yield a global resolution of 3.0 Å (GSFSC = 0.143).

#### HsFuzzy-Inturend-Rab23^N121I^

The data of *Hs*Fuzzy-Inturned-Rab23^N121I^ was processed accordingly (Fig. S3). After micrograph curation, 5960 micrographs with a CTF fit better than 5 Å were included for further data analysis. After initial blob picking, a Topaz wrapper was trained and a total of 220 thousand particles picked. Particle extraction was performed with a box size of 384 pixels (Foruier cropped to 96 pixels, resulting in a pixel size of 2.72 Å per pixel). Iterative rounds of 2D classification and selection were performed (Fig. S3). Selected classes were used for ab initio reconstruction in three classes followed by Heterogeneous and NU Refinements resulting in a one class of 113 thousand particles. 3D classification with forced hard classification enabled resulted in identification of a dimeric (Fuzzy-Inturned, 38 thousand particles) and a trimeric (Fuzzy-Inturned-Rab23^N121I^, 60 thousand particles) population, which were further refined in NU refinements at a box size of 384 pixels leading to final reconstructions of 3.6 Å and 3.4 Å resolution (GSFSC = 0.143), respectively.

All maps were subjected to unsupervised B-factor sharpening within cryoSPARC. No symmetry was applied during processing. All GSFSC curves, angular distribution plots and local resolution maps were generated with cryoSPARC (Supplementary Fig. 1-4). The local resolutions of refined maps (Fig. S1, S3) were estimated in cryoSPARC and analyzed in UCSF ChimeraX (*51*).

### Model building, Refinement and Validation

The structures of ctMon1-Ccz1 (PDB: 7QLA) (*12*) and Ypt7 (PDB: 5LDD) (*14*), the human CPLANE complex (PDB: 7Q3D) (*21*) and the AlphaFold prediction of Rab23 (*52*) were manually fitted as rigid bodies into maps using UCSF ChimeraX and used as a starting model. Subsequently, iterative rounds of real space refinement in PHENIX (*53*) and manual adjustments in WinCoot 0.9.8.92 (*54*) were performed. Model validation was done using MolProbity (*55*) in PHENIX. Models and maps were visualized and figures were prepared in UCSF ChimeraX. Model refinement and validation statistics are provided in Supplementary Table 1.

## Acknowledgments

We thank Ann-Marie Lawrence-Dörner and Bianca Berkenfeld for technical assistance, and the members of the Kümmel Lab for constructive feedback. We are grateful to Christian Ungermann and Lars Langemeyer for insightful discussions and to Francis Barr for providing plasmids encoding Fuzzy, Inturned, Rab23 and Rsg1. The template clone Flag-ciBAR1 was a gift from Ken-Ichi Takemaru (Addgene plasmid # 200440). This work was supported by the German Research Foundation (DFG) through the grants SFB1557-P10 (DK) and SFB1557-P11 (AM). JHS was supported by the Friedrich-Ebert Foundation. ML acknowledges funding from the European Research Council (ERC) under the European Union’s Horizon 2020 research and innovation program (grant agreement no. 101045340).

## Author contributions

Conceptualization: SW, DK Methodology: DJ, DDI, ML Investigation: SW, JT, JHS, AL, SA Visualization: SW, JT, DK Supervision: SVW, ML, AM, DK Funding acquisition: AM, DK Writing—original draft: SW, DK Writing—review & editing: SW, JHS, SVW, AM, DK

## Data and materials availability

All data needed to evaluate the conclusions of the paper are present in the paper and the Supplementary Materials. The model coordinates corresponding cryo-EM maps will be deposited in the Protein Data Bank and the Electron Microscopy databank.

