## Supplementary Information for "Mechanistic Plasticity of the RabGEFs Mon1-Ccz1 and Fuzzy-Inturned"

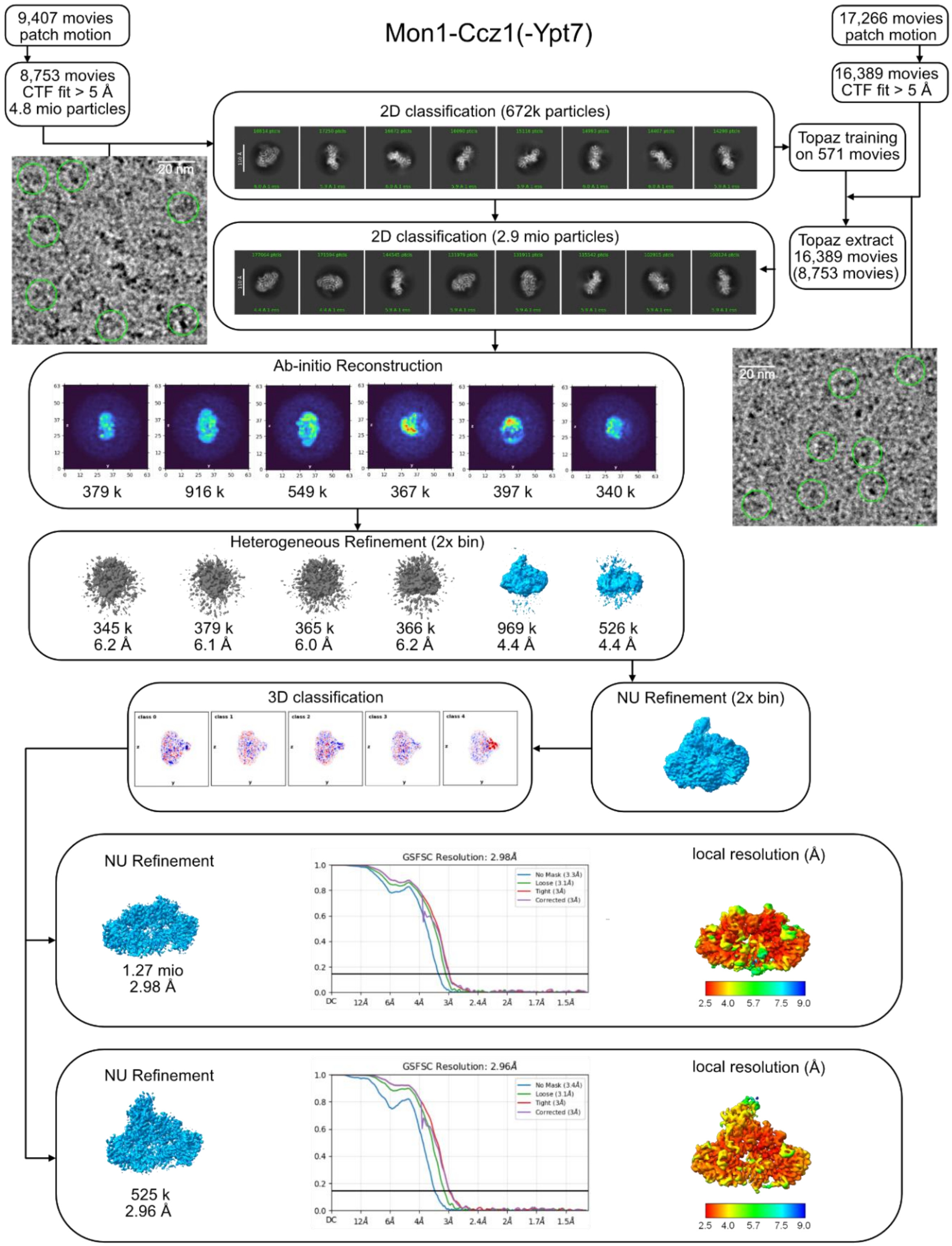


**Fig. S1.**
Cryo-EM data processing workflow for *Ct*Mon1-Ccz1-Ypt7 and *Ct*Mon1-Ccz1.

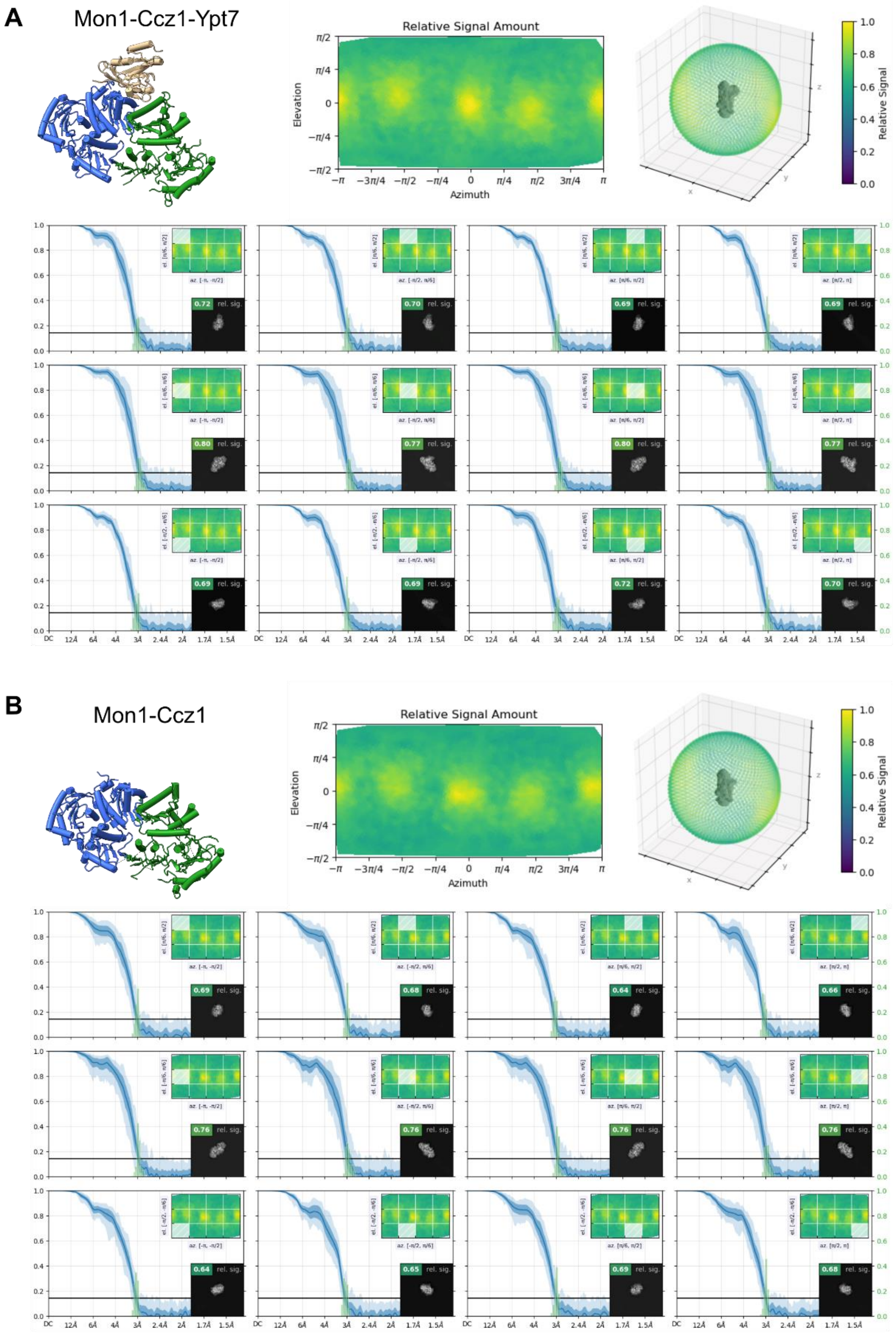


**Fig. S2.**
Orientation diagnostics for *Ct*Mon1-Ccz1-Ypt7 (A) and *Ct*Mon1-Ccz1 (B) cryo-EM data.

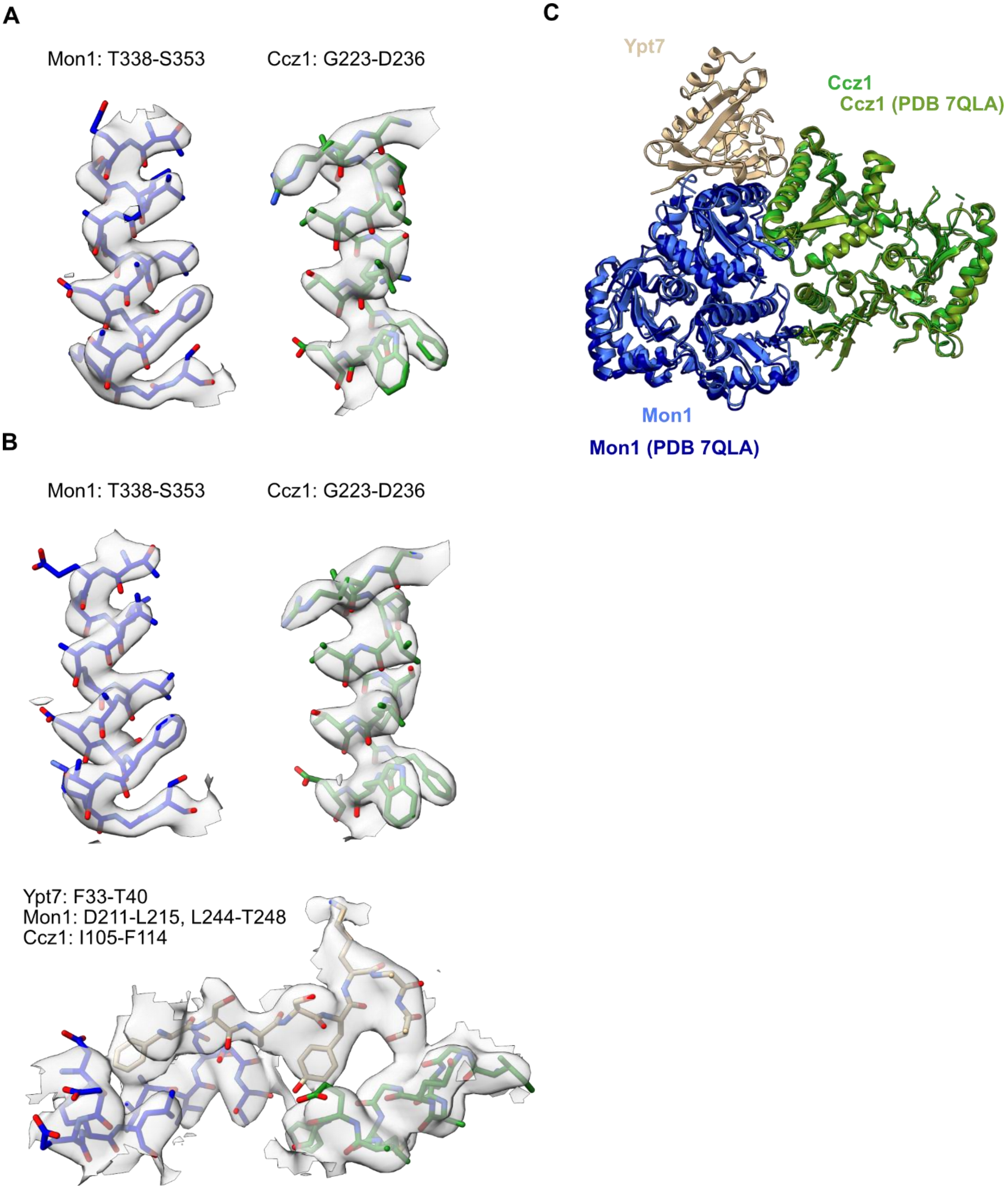


**Fig. S3.**
Model/map fit of selected areas of the (**A**) *Ct*Mon1-Ccz1 and (**B**) *Ct*Mon1-Ccz1-Ypt7 complexes. (**C**) Superposition of *Ct*Mon1-Ccz1$^{\Delta L}$-Ypt7 and *Ct*Mon1$^{\Delta N}$-Ccz1$^{\Delta L}$ (PDB ID 7QLA).

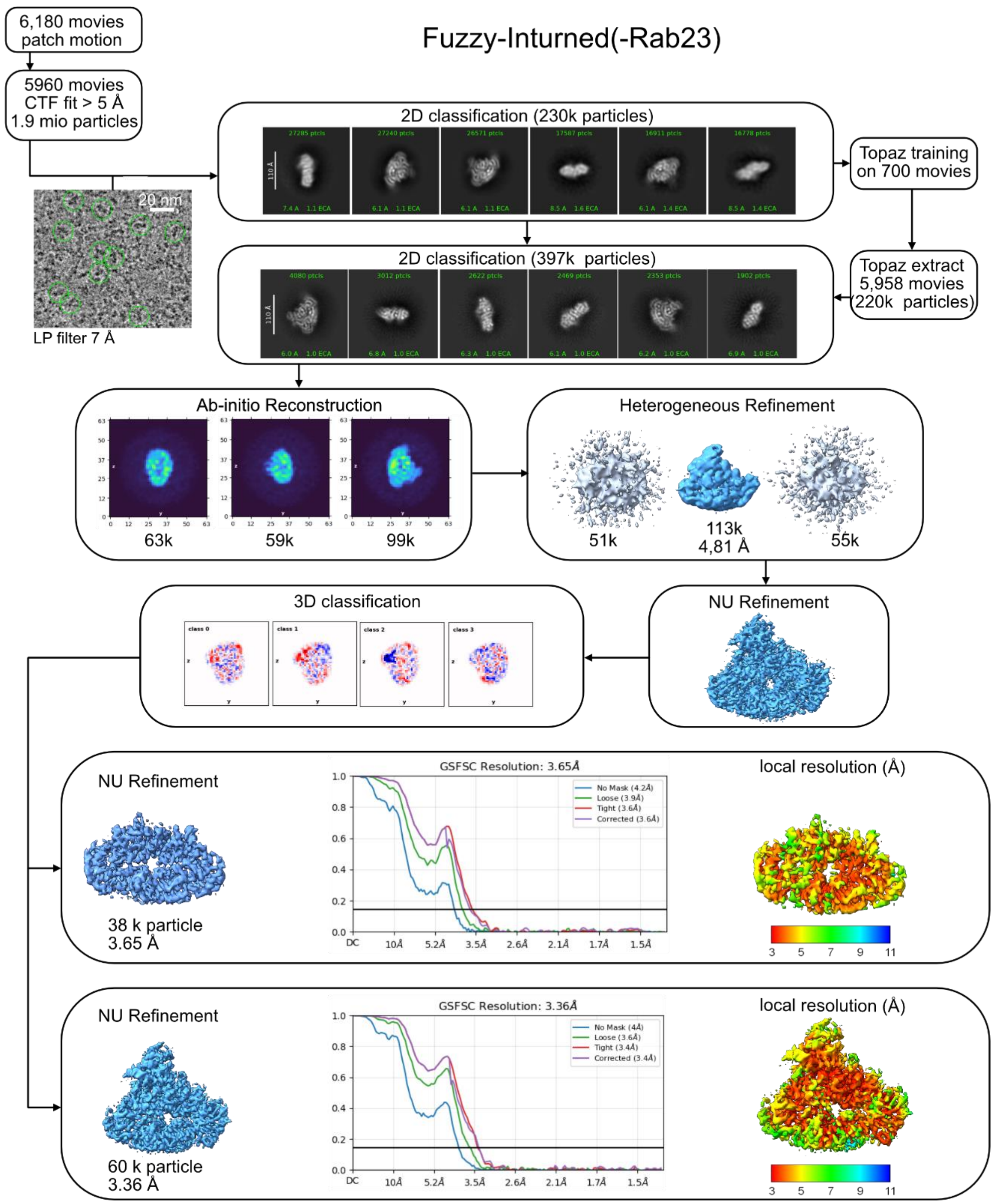


**Fig. S4.**

Cryo-EM data processing workflow for *Hs*Fuzzy-Inturned and *Hs*Fuzzy-Inturned-Rab23.

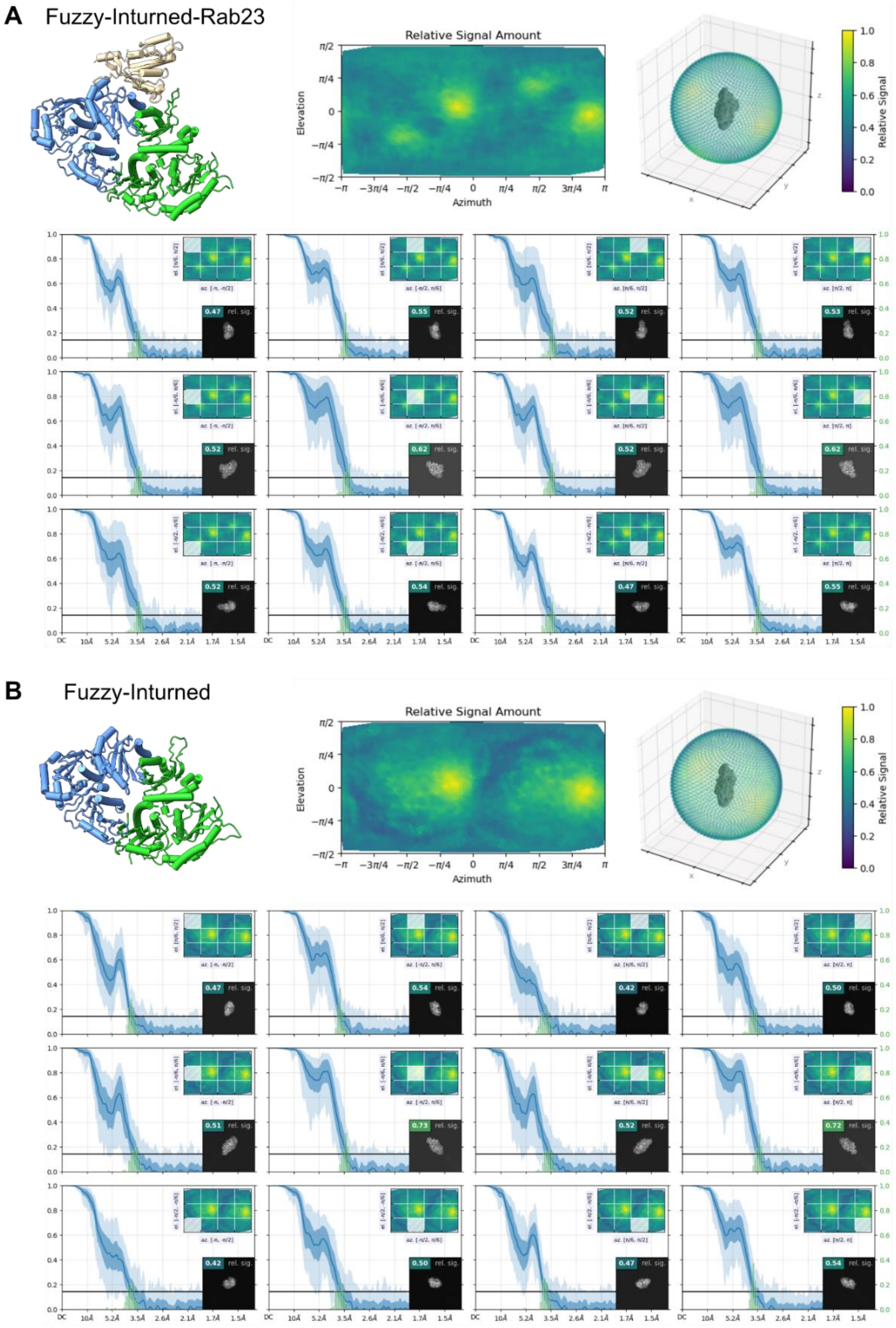


**Fig. S5.**
Orientation diagnostics for (**A**) *Hs*Fuzzy-Inturned-Rab23 and (**B**) *Hs*Fuzzy-Inturned cryo-EM data.

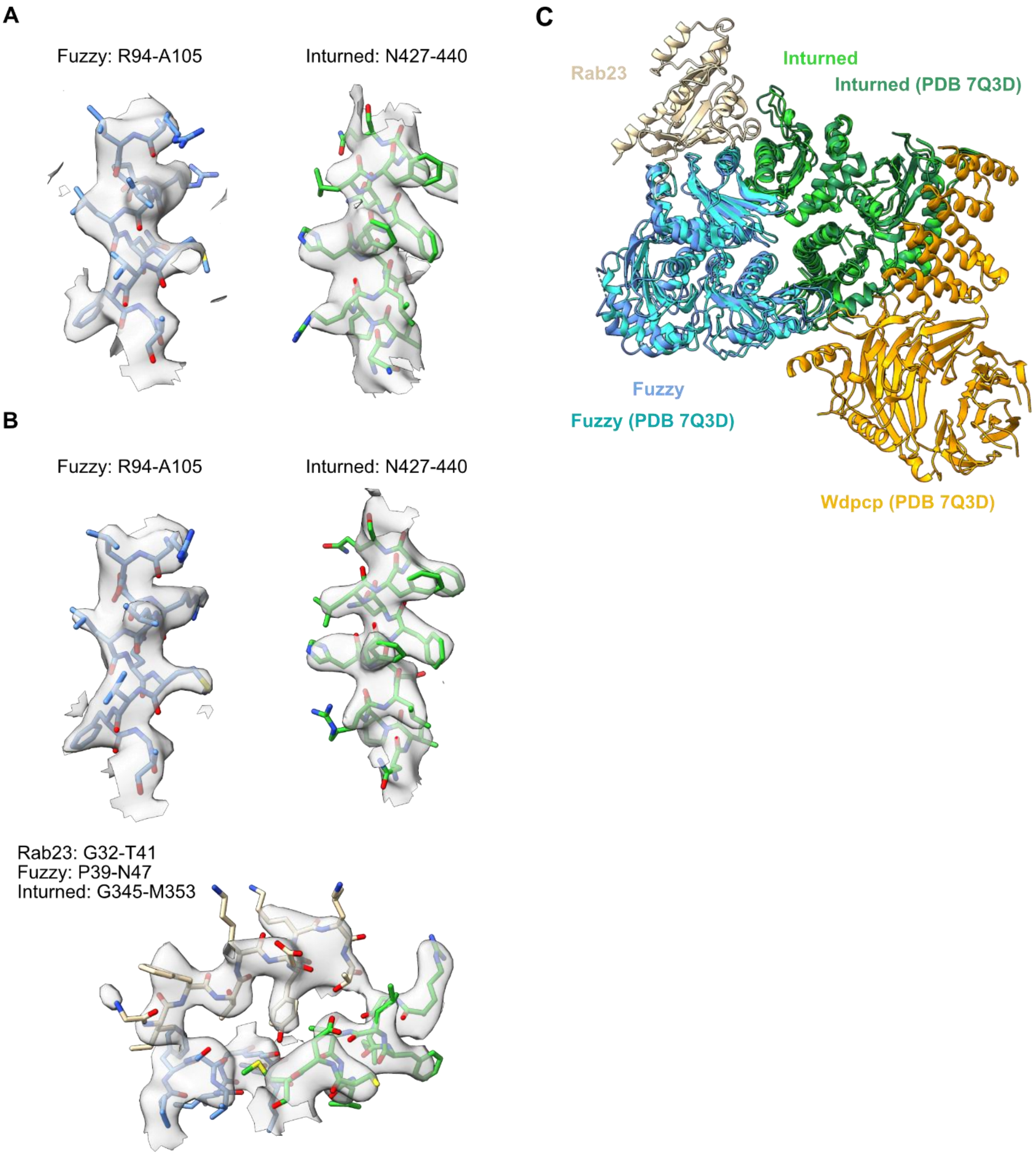


**Fig. S6.**
Model/map fit of selected areas of the (**A**) *Hs*Fuzzy-Inturned and (**B**) *Hs*Fuzzy-Inturned-Rab23 complexes. (**C**) Superposition of *Hs*Fuzzy-Inturned-Rab23 and the human CPLANE complex (PDB ID 7Q3D).

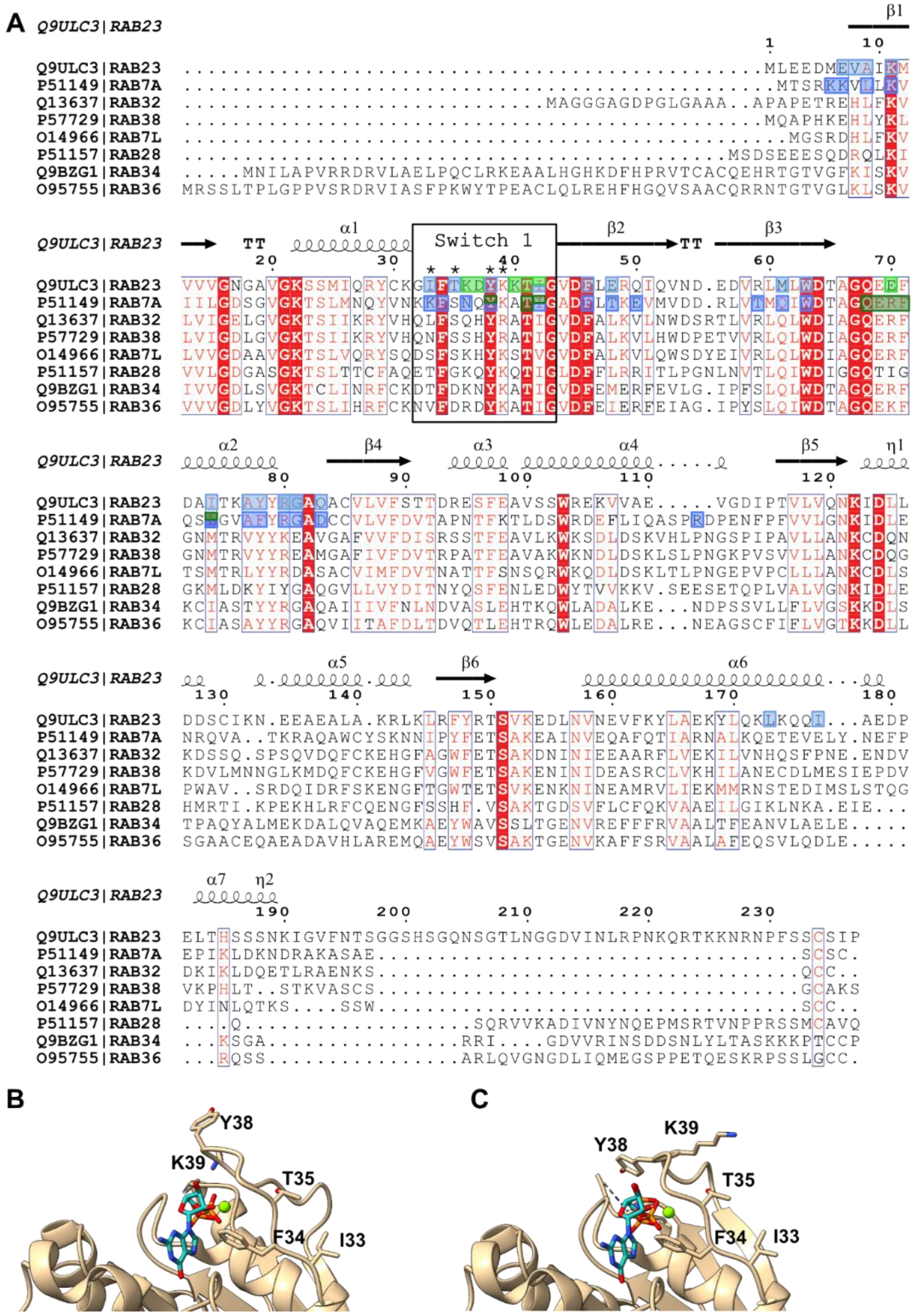


**Fig. S7.**
(**A**) Amino acid sequence alignment of human Rab GTPases containing FxxxYK/R motifs in their switch I. Secondary structure elements and sequence numbering are inferred from hsRab23. Residues mutated for GEF activity assays are highlighted by asterisks. Residues contacting GEF complexes are marked in blue (Mon1/Fuzzy) and green (Ccz1/Inturned). Note: Rab7 N35 differs from Ypt7 that was used for structural investigation and carries an Ala at that position. Crystal structures of hsRab23 in complex with (**B**) GDP (PDB ID 8YL3) and (**C**) the GTP analog GMPPNP (PDB ID 8YIM).

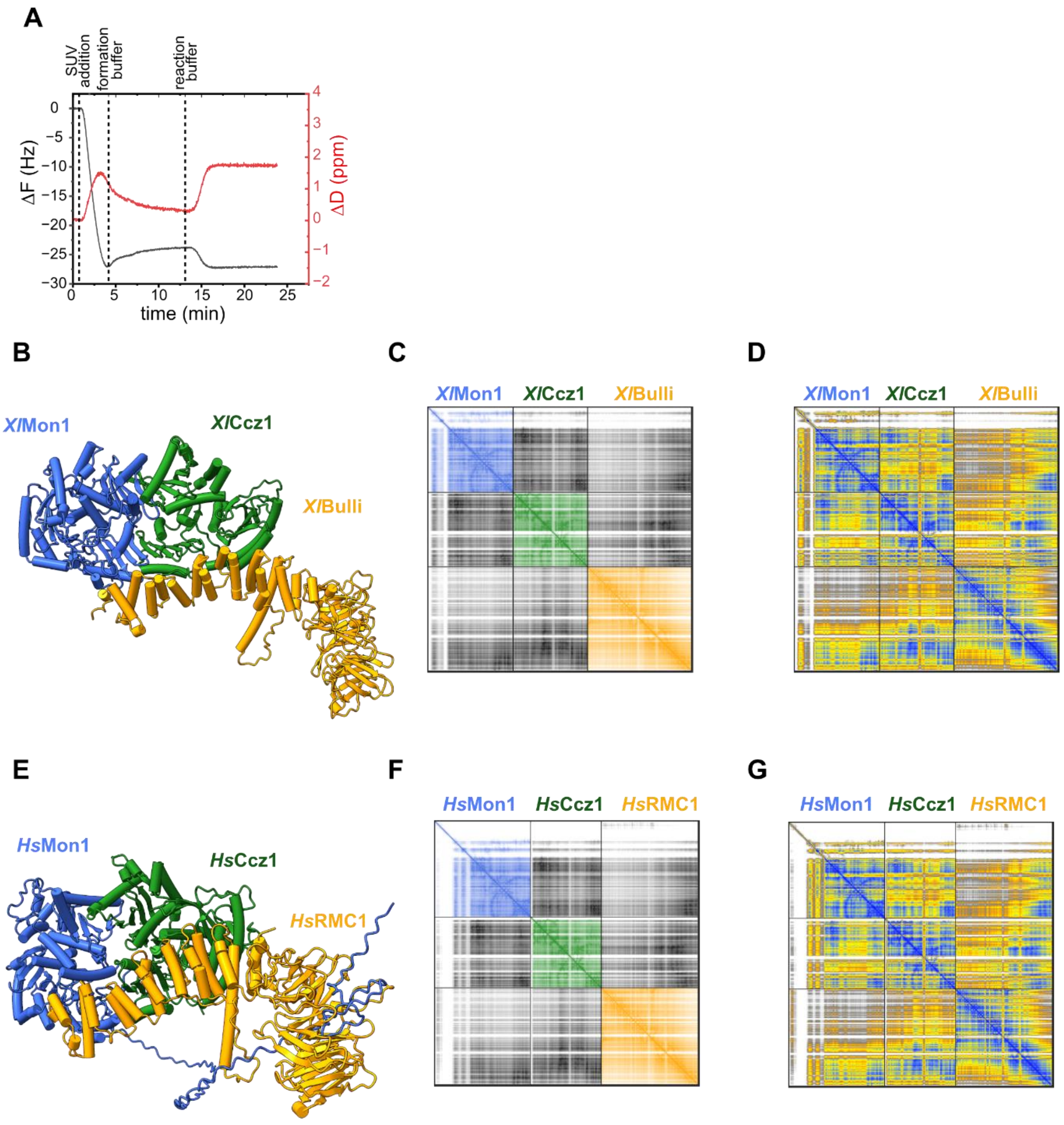


**Fig. S8.**
(**A**) Exemplary plot for frequency (black, deviation from resonance frequency, Δ/Hz) and dissipation (red, dissipation changes (ΔD)) shifts during bilayer formation for QCM-D measurements. Membrane composition: 74 mol% DOPC, 18 mol% DOPE, 2 mol% PI3P (18:1), 1 mol% PI3,5P2 (18:1) and 5 mol% PS. (**B**) Alphafold3 prediction of xlMon1-Ccz1-RMC1 with pTM = 0.81 and ipTM = 0.80 scores. (**C**) pAE plot for B colored by chain. (**D**) pAE plot for B colored by pLDDT. (**E**) Alphafold3 prediction of hsMon1-Ccz1-RMC1 with pTM = 0.78 and ipTM = 0.76 scores. (**F**) pAE plot for E colored by chain. (**G**) pAE plot for E colored by pLDDT.

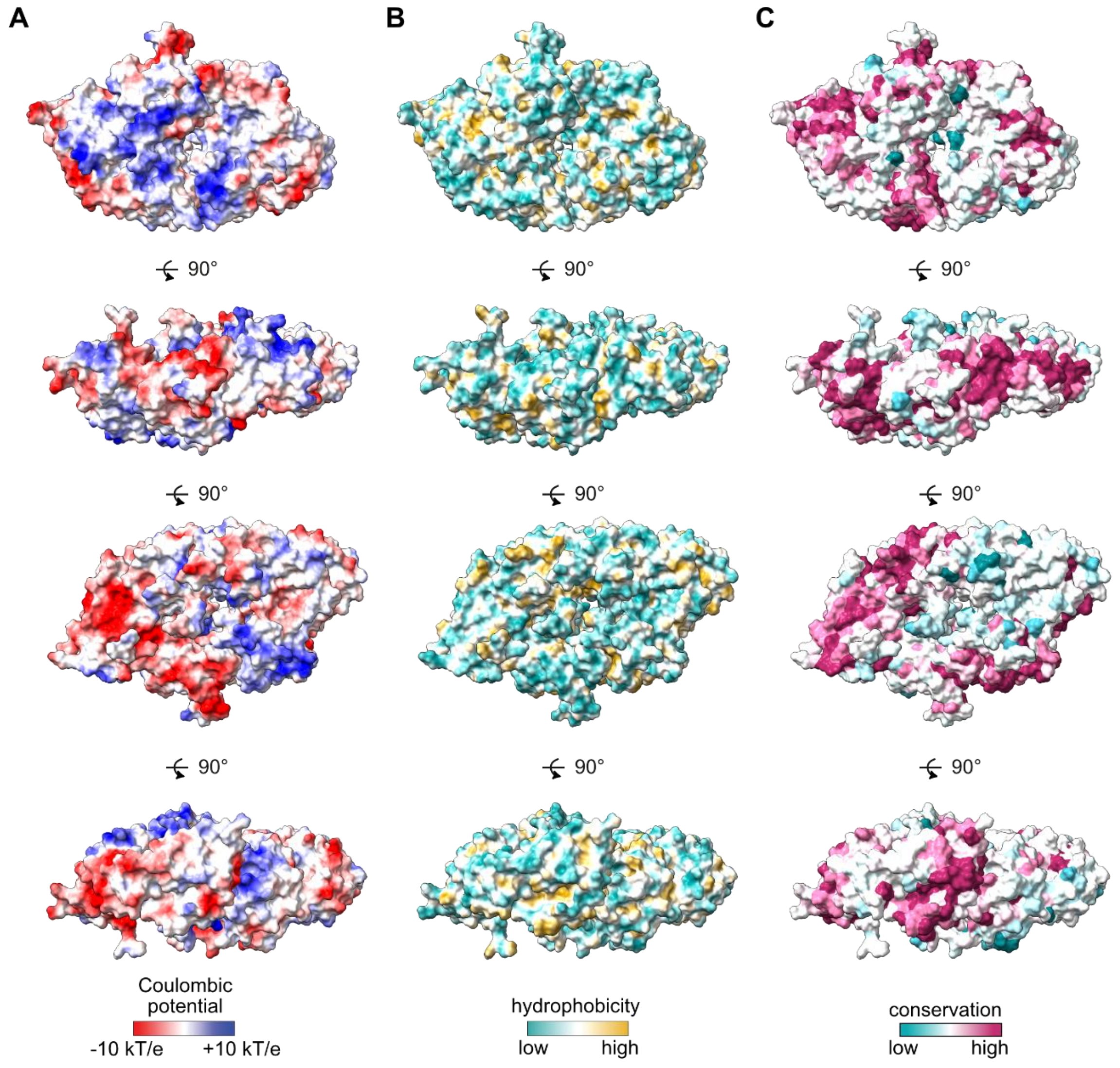


**Fig. S9.**
Surface representations of *Hs*Fuzzy-Inturned colored by (**A**) coulombic potential, (**B**) hydrophobicity and (**C**) conservation.

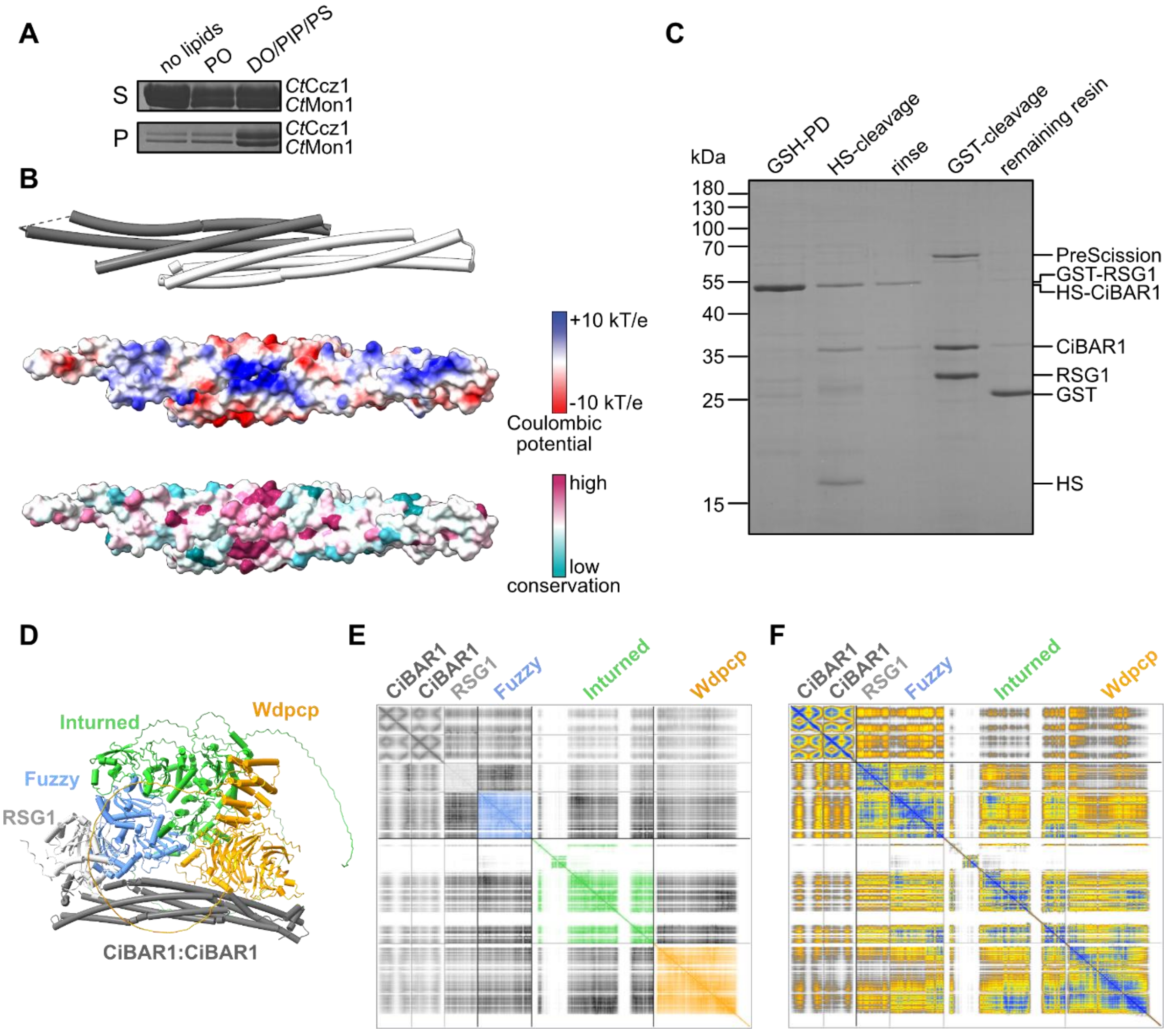


**Fig. S10.**
(**A**) The *Ct*Mon1-Ccz1 complex is recruited to membranes containing packing defects and charges in co-sedimentation assays. Lipids mixtures: PO: 81% POPC, 18% POPE, 1% DPPE-Atto 565 and DO-PIP-PS:73% DOPC, 18% DOPE, 1% DPPE-Atto 565, 2% PI3P, 1% PI3,5P2, 5% PS. (**B**) Crystal structure of the homodimeric BAR domain protein CiBAR1 (PDB ID: 8CEG) with surface representations of the coulombic potential and conservation. (**C**) Co-purification of His-SUMO (HS) tagged CiBAR1 and Glutathion-S-Transferase (GST) tagged Rsg1. Both proteins were co-purified on glutathione resin (GSH-PD). The HS tag cleaved off CiBAR1 by incubation with SUMO protease (HS-cleavage) and protease and tag rinsed away (rinse). The complex was finally eluted from the resin by cleaving the GST-tag with PreScission protease (GST-cleavage). (**D**) Alphafold3 prediction of CiBAR1(dimer) in complex with CPLANE (Rsg1, Fuzzy, Inturned, and Wdpcp) with pTM=0.71 and ipTM=0.71 scores. (**E**) pAE plot of D colored by chain. (**F**) pAE plot of D colored by pLDDT scores.

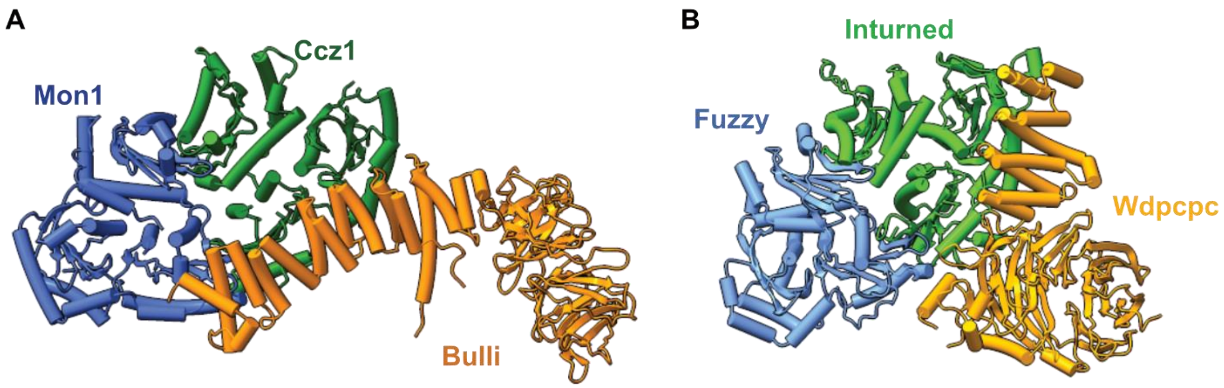


**Fig. S11.**
Side by side comparison of (**A**) *Dm*Mon1-Ccz1-Bulli (PDB ID: 8C7G) and (**B**) *Hs*Fuzzy-Inturned-Wdpcp (PDB ID 7Q3D). Despite high structural similarities between Bulli and Wdpcp, these subunits are integrated into the complex entirely different.

**Table S1. Cryo-EM data collection and refinement statistics**

| Data Collection | Mon1-Ccz1-Ypt7 | Mon1-Ccz1 | IntuFy-Rab23 | IntuFy |
|---|---|---|---|---|
| Microscope | FEI Glacios | | FEI Glacios | |
| Voltage (kV) | 200 | | 200 | |
| Camera | Falcon 4i | | Falcon 4i | |
| Energy filter | Selectris | | Selectris | |
| Pixel size | 0.680 | | 0.680 | |
| Micrographs | 26,673 (25,142 final) | | 6180 (5690 final) | |
| Particles | 524738, | 1,271,173 | 60,395 | 37,728 |
| Total electron does ($e^-$/ Å$^2$) | 50 | | 50 | |
| Defocus range (µm) | -0.8 to -2.5 | | -0.8 to -1.8 | |
| **Atomic model composition** | | | | |
| Symmetry imposed | C1 | C1 | C1 | C1 |
| Non-hydrogen (protein) atoms | 8249 | 6970 | 7423 | 6082 |
| Residues | 1051 | 891 | 939 | 772 |
| **Refinement (Phenix)** | | | | |
| RMSD bond length (Å) | 0.003 | 0.003 | 0.004 | 0.004 |
| RMSD angle (°) | 0.624 | 0.682 | 0.792 | 0.829 |
| Model to map fit, CC mask | 0.78 | 0.82 | 0.77 | 0.74 |
| Model to map fit, CC box | 0.70 | 0.81 | 0.82 | 0.82 |
| Resolution (FSC @ 0.143, Å) | 3.0 | 3.0 | 3.4 | 3.7 |
| B-factor (min/ max/ mean, Å$^2$) | 53/196/108 | 86/242/141 | 47/218/124 | 109/233/150 |
| **Validation** | | | | |
| Clash score | 16.73 | 9.93 | 21.85 | 16.0 |
| Ramachandran outliers (%) | 0.00 | 0.00 | 0.00 | 0.00 |
| Ramachandran allowed (%) | 4.42 | 3.70 | 5.72 | 4.0 |
| Ramachandran favoured (%) | 95.58 | 96.30 | 94.28 | 96.0 |
| Molprobity score | 2.44 | 1.77 | 2.22 | 1.98 |
| **PDB accession** | XXXX | XXXX | XXXX | XXXX |
| **EMDB accession** | XXXX | XXXX | XXXX | XXXX |
